# Retinal and callosal activity-dependent chandelier cell elimination shapes binocularity in primary visual cortex

**DOI:** 10.1101/612358

**Authors:** Bor-Shuen Wang, Maria Sol Bernardez Sarria, Miao He, Michael C Crair, Z. Josh Huang

## Abstract

In mammalian primary visual cortex (V1), integration of the left and right visual scene into a binocular percept derives from convergent ipsi- and contralateral geniculocortical inputs and trans-callosal projections between the two hemispheres. However, the underlying developmental mechanisms remain incompletely understood. Using genetic methods in mice we found that during the days before eye-opening, retinal and callosal activity drives massive apoptosis of GABAergic chandelier cells (ChCs) in the binocular region of V1. Blockade of ChC elimination resulted in a contralateral-dominated V1 and deficient binocular vision. As activity patterns within and between retinas prior to vision convey organization of the visual field, their regulation of ChC density through the trans-callosal pathway may prime a nascent binocular territory for subsequent experience-driven tuning during the post-vision critical period.

**One Sentence Summary:** Prior to eye opening the developing retina primes the visual cortex for binocular vision by adjusting the density of a cortical inhibitory neuron type.

## Main Text

In mammals with more front-oriented eyes, the seamless integration of the left and right visual scene into binocular vision is crucial for many visually guided behaviors. Information about the central visual field is relayed from the left and right temporal retinas separately to the lateral geniculate nuclei (LGN) and then converges in the lateral regions of V1, defining the binocular zone (BZ) (*1*). In addition, a set of V1 pyramidal neurons (PyNs) project to the contralateral BZ, forming a trans-callosal pathway that also contributes to binocular vision (e.g. up to 40-50% of ipsilateral eye responses in rodents (*2*-*4*)). The developmental assembly of binocular circuits from retina to V1 is not well understood. Pioneering studies demonstrated visual experience-dependent tuning of V1 binocular properties during a postnatal critical period (*5*), a neural plasticity principle that manifests across mammalian species (*6*). On the other hand, the spatial and topographic organization of retinal inputs reflect cardinal features of the visual world, including the central meridian bridging the left-right visual field. this information is conveyed along the visual pathway to V1 before eye opening as spatiotemporal patterns of spontaneous activity within and between the two eyes (*7*, *8*). How early retinal activity and the developing trans-callosal pathway shapes binocular properties in V1 is not known. In particular, the role of specific cortical GABAergic interneurons in this process has not been explored.

ChCs are among the most distinct cortical interneurons. They control PyN spiking at the axon initial segment (*9*). We labeled ChCs through their embryonic progenitors by tamoxifen administration to pregnant *Nkx2.1-CreEr;Ai14* mice (*10*) and examine their distribution across V1 and lateral secondary (V2L) visual areas (Fig. 1A-B, fig. S1). To demarcate the BZ, we injected AAV-GFP into the contralateral hemisphere and visualized callosal axons innervating the V1/V2L border, which was further identified by the expression pattern of type 2 muscarinic acetylcholine receptors (*11*) (Fig. 1A-B, fig. S1, see Methods). We found that in mature visual cortex (>P28), the density of layer 2 (L2) ChCs in the border region was half that in neighboring V1 or V2L (Fig. 1G,H). Notably, there was no such areal difference in Parvalbumin or Calretinin interneurons (fig. S1). Furthermore, reduced ChC density was not evident in other cortical areas receiving callosal projections, such as the S1-S2 and M1-S1 borders (fig. S2).

**Fig. 1.**
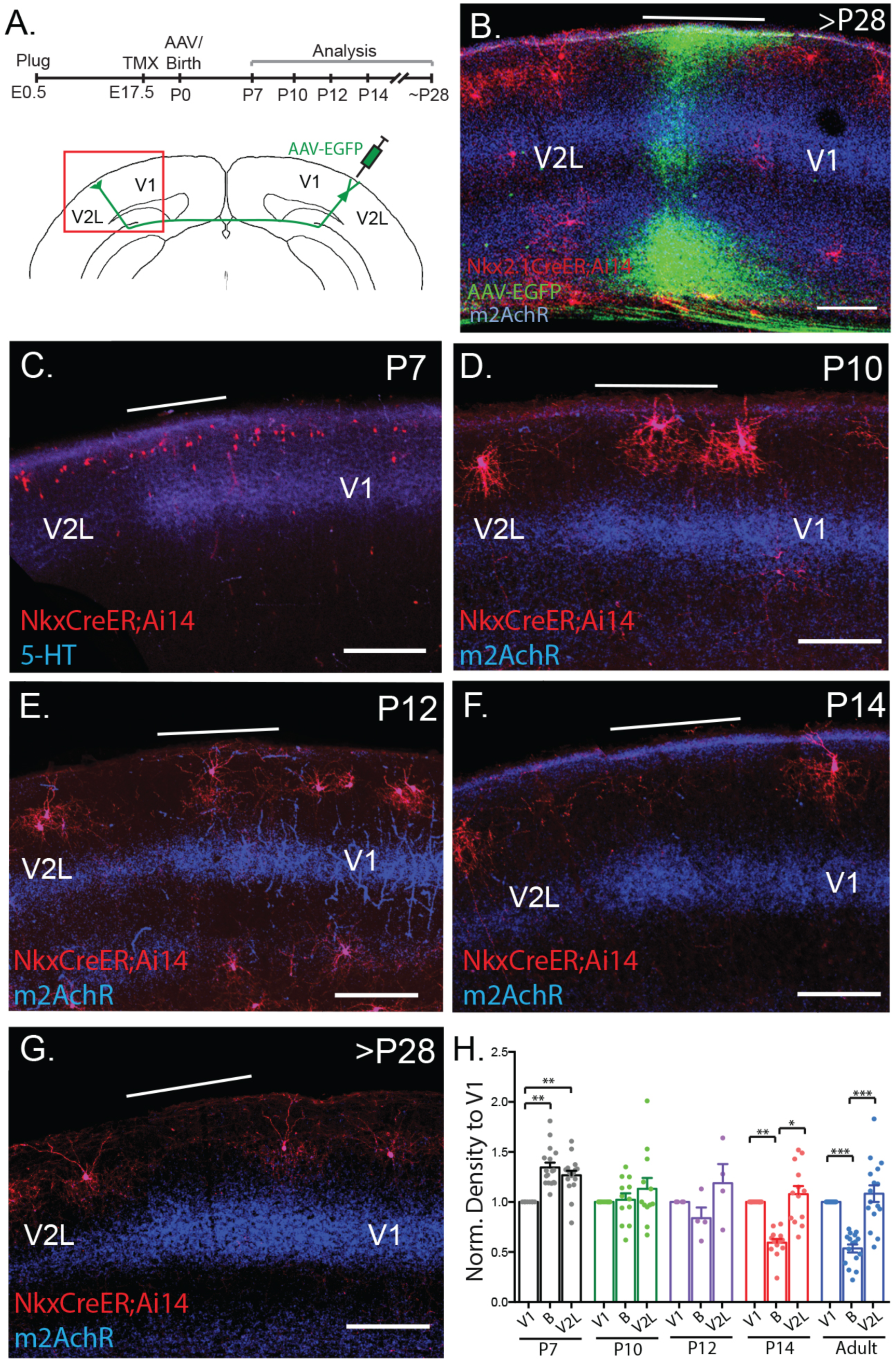
Massive ChC elimination at V1/V2L border during second postnatal week. (**A**) Schematic of experimental design. AAV-EGFP was injected into visual cortex at P0 and ChC density in contralateral cortex analyzed at indicted time points. (**B**) Example AAV-EGFP labeled callosal axon projection into contralateral V1/V2L border region where ChCs express RFP. (**C-G**) ChC distribution at P7 (n=17), P10 (n=12), P12 (n=4), P14 (n=13), P28 (n=16). White line marks the V1/V2L border. (**H**) Normalized ChC density to V1 across development. At P7, ChC density at the callosal axon recipient zone is initially higher than V1; ChCs are progressively eliminated between P7-P14, reaching adult-like density by P14. * p < 0.05, ** p < 0.01, *** p < 0.001. Scale bar: 200 um.

Following their generation from the medial ganglionic eminence (MGE) (*10*, *12*) and migration into the cortex, a large subset of young ChCs settle in L2 by the end of the first postnatal week and densely distributed across cortical areas including V1 and V2L (Fig. 1C). Between P7-P14, however, ChC density progressively and permanently decreased in the V1/V2 border region to half that of neighboring regions (Fig. 1C-H). Therefore, following an initial adjustment of MGE interneuron density during the first postnatal week, which is dependent on the level of glutamatergic input from nearby PyNs (*13*), ChC density specifically and dramatically decreased in the BZ of V1 in the second postnatal week.

To investigate the cellular mechanism of ChC elimination, we blocked apoptosis in ChCs by crossing *Nkx2.1-CreER;A14* mice to *Bak*^*-/-*^*Bax*^*f/-*^ conditional knockout mice (*14*). At P30, the ChC density in the BZ of these mice was comparable to that of nearby V1 and V2L (fig. S3D), and closely resembled the distribution pattern of WT P7 mice (before ChC elimination; fig. S3E). Interestingly, we observed two populations of ChCs in apoptosis-blocked mutants: one with normal morphology and another with a significantly smaller cell body and short neurites that were concentrated at the V1/V2L border (fig. S3F). These atrophic ChCs likely represent those that would have been eliminated during normal development, but remained due to apoptosis blockade.

Developing L2 ChC dendrites arborize mainly in L1 and likely receive local, thalamic, and contralateral callosal inputs (*15*). As the spatiotemporal pattern of L2 ChC apoptosis closely correlates with the invasion of callosal axons (*16*, *17*), we investigated the possible link between these two events. We used in utero electroporation at embryonic day 15.5 (E15.5) to label L2/3 callosal PyNs with EGFP (Fig. 2A). Callosal axons extended to the contralateral hemisphere and arborized densely in a restricted border region between V1 and V2L (Fig 2B,C). Co-electroporation of a potassium channel Kir2.1, which reduces neuronal activity, dramatically reduced the growth and invasion of callosal axons into contralateral cortex (Fig. 2D), consistent with previous findings (*16*, *17*). Importantly, reduced callosal axon innervation resulted in an increase in ChC density at the border (Fig. 2D,H,I), suggesting that callosal axon invasion promotes ChC elimination.

**Fig. 2.**
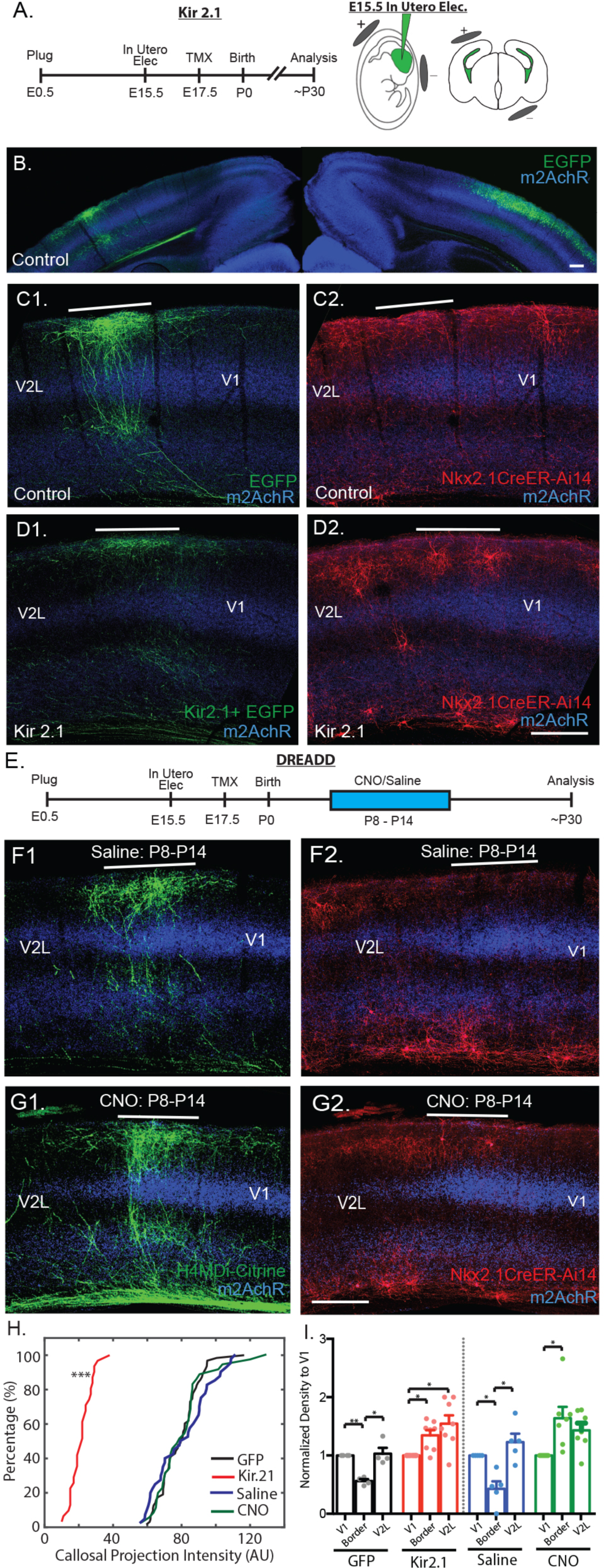
Trans-callosal axons and activity regulate ChC elimination at V1/V2L border. (**A**) Experimental design with CAG-Kir 2.1-EGFP or CAG-EGFP electroporation at E15.5 targeting layer 2/3 callosal PyNs in visual cortex and TMX labeling of ChCs at E17.5. (**B-C**) In a control animal, EGFP-labeled V1 L2/3 PyNs sent callosal axons innervating the contralateral V1/V2L border (**C1**) where few ChCs were present (**C2**). (**D**) Kir2.1 IUE reduced callosal axonal projection to V1/V2L border (**D1**), which prevented ChC elimination at border (**D2**). (**E**) Schematic of suppressing callosal activity with H4MDi-Citrine electroporation at E15.5 and daily CNO i.p. injections between P8-P14. (**F-G**) Silencing callosal neuron activity between P8-P14 did not affect callosal projection into contralateral cortex (**F1**,**G1**), but prevented ChC elimination at the border (**F2**,**G2**). (**H**) Cumulative distribution of callosal axon intensity at the contralateral V1/V2L border showing lack of projections in Kir2.1-expressing animals (p < 0.001, KS test), but normal projection intensity in H4MDi-treated animals. (**I**) Normalized ChC density relative to V1 showing rescue of ChCs at the border region in Kir2.1-expressing animals (n=8) compared to GFP controls (n=4), and in H4MDi-expressing animals treated with CNO (n=7) compared with saline controls (n=5). * p<0.05, ** p<0.01, *** p<0.001. Scale Bar: 200um.

To examine the role of callosal activity in regulating ChC survival, we expressed Hm4Di (inhibitory DREADD receptor) (*18*) in L2/3 PyNs via electroporation at E15.5 (Fig. 2E,F) or virally via cortical injection at P0-1 (fig. S4). Application of clozapine-N-oxide (CNO) suppressed firing of V1 PyNs both in vitro (fig. S5) and in vivo (fig. S6). Daily application of CNO from P8-P14 prevented ChC death at P28, leading to significantly higher ChC density in the contralateral BZ compared to saline and CNO alone controls (Fig. 2F-I; fig. S7), without affecting callosal projection patterns (Fig 2G,H). Suppression of PyN activity via CNO led to reduced interhemispheric activity correlations without affecting within-hemisphere correlations (2-way ANOVA between treatment and visual subregion; F=4.44, p=0.02; fig. S6) Together these results suggest that ChC elimination at the V1/V2L border is regulated by callosal PyN activity, likely through coordinating callosal neuron activity across the two hemispheres.

The precision of callosal projections to the contralateral V1/V2L border region is shaped by retinal inputs during a perinatal critical period (*19*). Perinatal monocular enucleation (ME) in rats induces broadened and ectopic callosal projections into V1 (*19*). We found that P0 ME in mice also resulted in broadened callosal terminations in contralateral V1/V2L (Fig. 3A-D). Importantly, this led to a correspondingly expanded reduction in ChC density (Fig. 3E), indicating that invading callosal axons were sufficient to eliminate ChCs even in ectopic locations. Therefore, early retinal inputs not only play a crucial role in shaping trans-callosal axon projections to the BZ but also regulate ChC survival at the V1/V2L border.

**Fig. 3.**
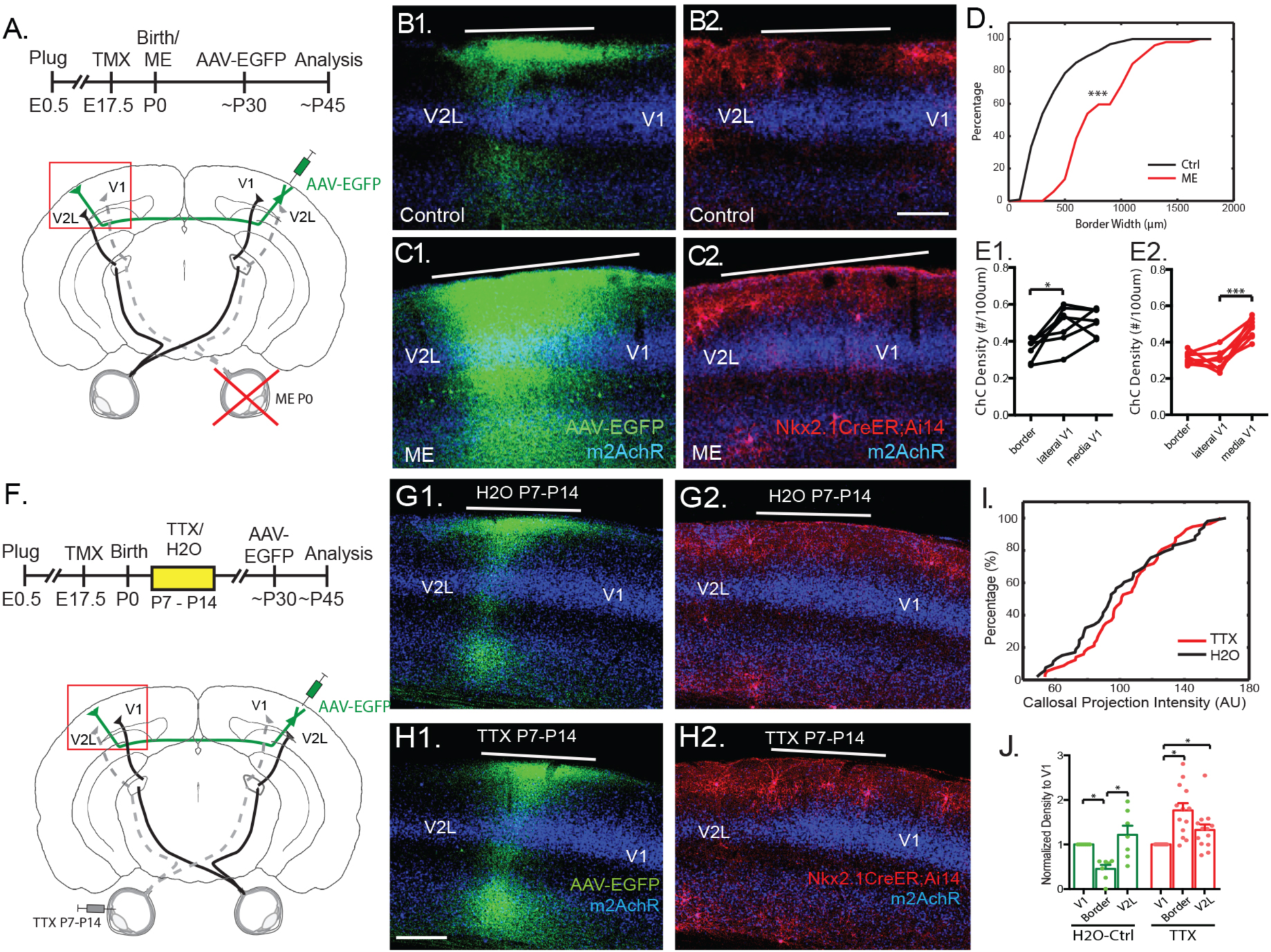
Ectopic callosal axon projections promotes ChC elimination within V1 (**A**) Schematic of experimental design. (**B**) Control animal with normal callosal projection to V1/V2L border (**B1**) and lack of ChCs at border (**B2**). (**C**) ME animal with broad and ectopic callosal projections into V1 (**C1**), and a reduction in ChC density within V1 (**C2**). (**D**) Widening of callosal projection axon innervations into V1 in ME animals at P0 (p<0.001, KS test). (**E**) Reduction in ChC density within lateral portion of V1 in ME animals due to widening of callosal recipient zone (Control: n=7; ME: n=8). (**F**) Schematic of manipulating retinal activity with intravitreal TTX injection. (**G**) In control H2O intravitreal injected animal, EGFP-labeled callosal projection to the contralateral V1/V2L border (**G1**) with few ChCs (**G2**). (**H**) Example of TTX-injected animals with EGFP-labeled callosal axons (**H1**) and ChCs remaining at the border (**H2**). (**I**) Similar callosal projection intensity at V1/V2L border between TTX and H2O treated controls. (**J**) Normalized ChC density relative to V1 showing rescue of ChCs at the border after TTX injection (Control: n=7; TTX: n=13). * p < 0.05, ** p < 0.01, *** p < 0.001. Scale Bar: 100um.

Prior to eye opening, developing retinal ganglion cells generate spontaneous and correlated waves of activity that sweep across the retina (*8*, *20*) (*21*, *22*) and are enhanced by light just prior to eye opening (*23*). Retinal waves preferentially initiate in the ventral-temporal retina, which relays information about the binocular visual field to the V1/V2L border region of cortex where trans-callosal axons innervate (*7*). To examine how retinal activity might influence the bilateral coordination of cortical activity and ChC survival, we enucleated one eye at P6, prior to ChC elimination in the BZ. While callosal projections were not affected (fig. S8A,B), consistent with previous studies (*19*), ChCs were spared from apoptosis at the V1/V2L border (fig. S8B,C). To further explore the role of retinal activity in this process, we unilaterally blocked retinal activity between P7-P14 through intravitreal injection of the potent sodium channel blocker, tetrodotoxin (TTX) (Fig. 3F). While callosal projections were not affected by this manipulation (Fig. 3G-I), ChC elimination at the V1/V2L border was compromised (Fig. 3J). Notably, bilateral blockade of retinal activity between P7-P14 also reduced ChC elimination at the V1/V2L border (fig. S9), suggesting that the overall level and coordination of retinal activity promotes ChC elimination in the BZ. Together, these results suggest that spontaneous activity from the retina plays a key role in regulating ChC density in the BZ by coordinating callosal neuron activity across the two hemispheres.

To investigate the functional impact of a developmental reduction of ChC density at the V1/V2L border, we examined neuronal response properties in V1 binocular neurons in animals with excess ChCs following suppression of callosal activity during the second postnatal week (Fig. 2G, fig. S4). As expected, control mice that underwent a saline treatment between P8-P14 showed a slight contralateral bias in the BZ (Ocular Dominance Index (ODI) of 0.03 ± 0.04; (*24*, *25*)). However, the ODIs of mice with excess ChCs were shifted toward the contralateral eye (mean ODI = 0.21 ± 0.03), and their contralateral/ipsilateral response ratios (C/I ratio) were significantly higher than control mice (Fig. 4A-C). We then analyzed the maximum response rate of cortical units following separate stimulation of each eye to further explore changes in contra- and/or ipsi-lateral responses. We observed a consistent reduction of ipsilateral, but not contralateral responses in mice with excess ChCs (Fig. 4D), and the number of ChCs at the border positively correlated with their ODI shift (Fig. 4E; fig. S10D). Monocular response properties, such as orientation selectivity, direction selectivity, spatial frequency preference, and binocular matching of orientation preference were normal in ChC-excess animals (fig. S10). Therefore, the persistence of excess ChCs at the V1/V2L border disrupts the binocularity of V1 neurons by suppressing responses to the ipsilateral eye.

**Fig. 4.**
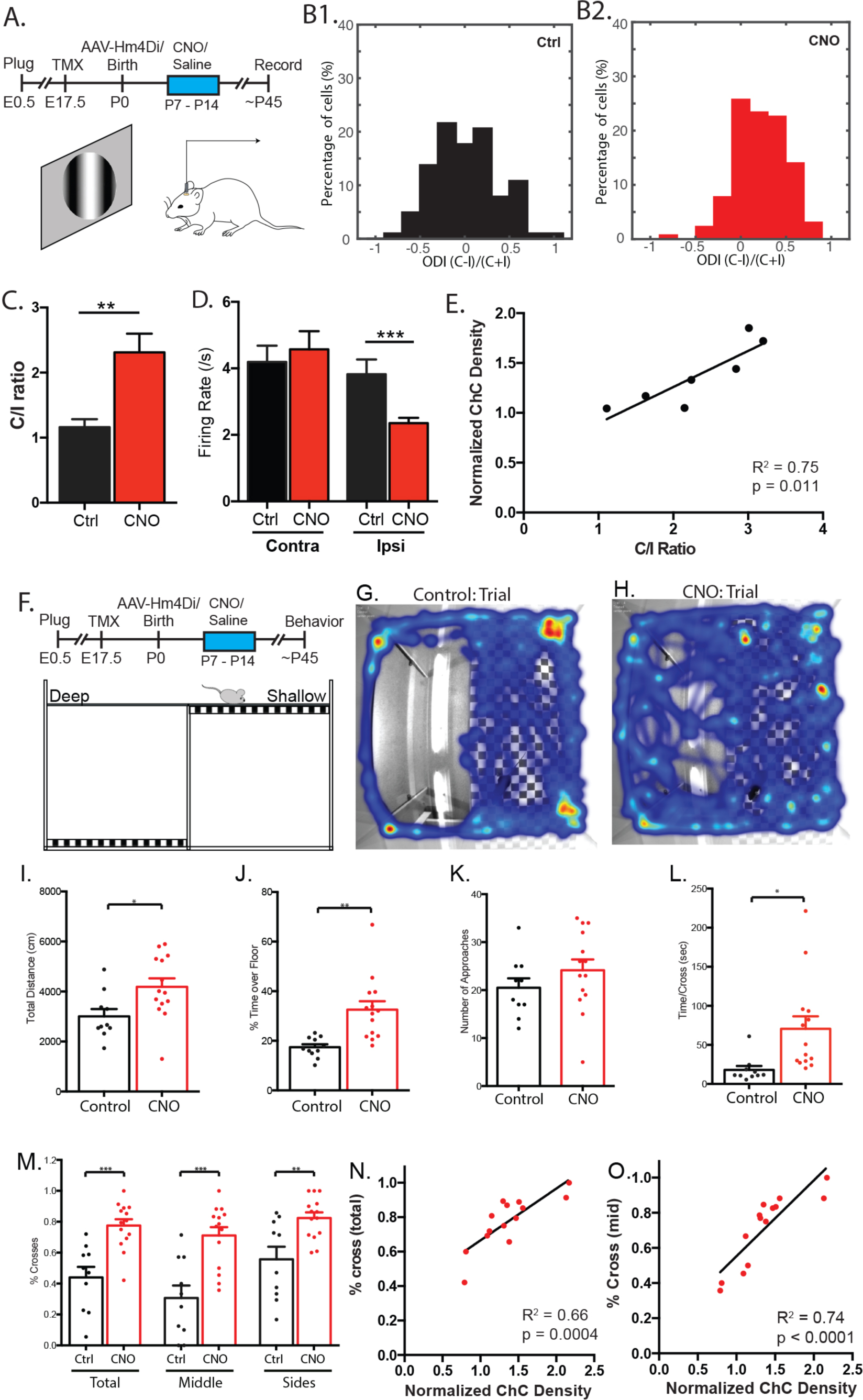
Excess ChCs at V1-V2L border perturbs binocularity and depth perception. (**A**) Experimental designing for assaying response properties in ChC-excess V1 using extracellular recording in lightly anesthetized mice. (**B**) Distribution of ODI in control (**B1**, n=102 cells, 5 animals) and CNO-treated (**B2**, n=128 cells, 7 animals) animals. (**C**) contralateral/ipsilateral response ratio (C/I ratio) showing a shift in ocular dominance in CNO-treated animals (p<0.01). (**D**) Weakened ipsilateral eye responsiveness in CNO-treated animals (n=128 cells, 7 animals) than controls (n=102 cells, 5 animals). (**E**) ChCs density at border region is positively correlated with C/I ratio in CNO-treated animals. (**F**) Schematic of visual cliff test. (**G-H**) Example trial movement tracking of control (G) and CNO-treated (H) animals. CNO-treated animals travels more distances (**I**) and spend more time over at the floor side (J; 32.5 ± 3.4%; n = 14) than the control (17.5 ± 1.3%; n = 10). (**K**) Both CNO-treated and control animals make similar numbers of approaches to cross the “cliff” (Control: 20.5 ± 1.9; CNO: 24.1 ± 2.3; p = 0.26). (**L**) The average time spent on the cliff side of the box per crossing was almost 4-fold higher in CNO-treated animals (Control: 17.9 ± 5.1 s; KO: 70.5 ± 16.0 s). (**M**) CNO-treated animals cross the “visual cliff” more often than control (Total: Control: 44.0 ± 6.8%; CNO: 77.6 ± 4.0% of approaches, n = 14; Middle: Control: 30.7 ± 8.0%; CNO: 71.1 ± 5.4%). (**N-O**) ChCs density at border region is highly correlated with the percentage of center crossing in CNO-treated animals. * p < 0.05, ** p < 0.01, *** p < 0.001.

Changes in binocular properties due to excess ChCs may impact visually guided behavior. To test stereopsis, we used a visual cliff test in which a high-contrast checkerboard is placed at two different heights below an open field arena with clear floors. This behavioral paradigm creates the visual illusion of a cliff (Fig. 4F) and mice with normal binocular vision tend to remain in the apparent shallow side (*26*). ChC-excess mice frequently crossed the visual cliff without caution or hesitation (video, Fig. 4H), while control mice behaved as expected, detecting the cliff and usually remaining in the shallow side of the arena (video, Fig. 4G). ChC-excess mice also spent double the time in the cliff region relative to controls (Fig. 4I,J). Although both groups approached the cliff at a similar frequency, ChC-excess mice crossed it significantly more frequently (Fig 4K,M). Mice may use the arena walls to explore and cross the visual cliff. We analyzed crosses at the side walls and center separately, and found ChC-excess mice crossed the cliff significantly more than controls independent of arena walls (Fig 4M). The average time spent on the cliff side per crossing was almost 4-fold higher in ChC-excess mice (Fig 4L). Interestingly, the density of ChCs at the border region was tightly correlated with the percentage of center crossing in ChC-excess animals (Fig 4N-O), whereas the distance travelled and duration spent were weakly correlated (Fig S11). Together, these results suggest that excess ChCs at the V1/V2L border leads to deficient binocular vision.

The assembly of eye-to-brain connections for binocular vision involves sequential steps that recruit developmental programs guiding axon growth and their topographic organization to activity and subsequently experience dependent refinement of synaptic connections (*1*). Here we have discovered a crucial step bridging these events: retinal activity before eye opening, which conveys information about the binocular visual field that influences the developing trans-callosal pathway, regulates the density of a powerful inhibitory interneuron type in the binocular region of V1 - a necessary step for subsequent experience-dependent development of binocular properties and binocular vision (fig. S12). As the contralateral eye dominates cortical responses at eye opening (*27*, *28*), ChC elimination may prime the nascent V1 for visual experience-driven strengthening of ipsilateral responses through either thalamo-cortical and/or trans-callosal connections (*3*) through ocular dominance plasticity during the post-vision critical period (*6*). Reduction of ChC-mediated inhibition in the BZ might also secure fast and efficient trans-callosal (*29*-*31*) communication, which is thought to link left and right V1 neurons with similar retinotopic and receptive field properties for achieving seamless integration of visual scenes, stereopsis, and motion tracking (*2*, *32*-*35*).

The mechanism of retinal and callosal activity regulation of ChC apoptosis at V1 BZ remains to be elucidated. It is possible that young ChCs initially innervate both non-callosal and callosal PyNs, while the latter establish bilateral reciprocal excitatory connections. As GABAergic transmission may be depolarizing at this postnatal period (*36*, *37*), ChCs innervating callosal PyNs might promote their firing, forming a trans-callosal loop that is further driven by coordinated bilateral retinal inputs. Such a transient over-excited network may drive the elimination of mis-wired ChCs through apoptosis, thereby shaping a fast trans-callosal network by preventing ChC-mediated inhibition at later stages. Young ChCs in other cortical areas receiving callosal inputs may also be wired into a similar excitatory loop, but the bilateral correlation of peripheral inputs to other areas (e.g. barrel, auditory cortex) (*38*, *39*) may not be synchronous enough to produce the level of over-excitation that eliminates ChCs in V1. Although early PV and SST cells may also depolarize callosal PyNs, their targeted synapses at the soma and dendrite might be less effective in promoting PyN firing. Our results suggest a mechanism whereby the organization of the peripheral sense organ, which reflects cardinal features of the physical world, regulates the developmental integration of cortical inhibitory interneurons and primes subsequent experience-dependent tuning of sensory perception.

## Acknowledgments

We thank Xu An for providing software coding for in-vivo electrophysiology data acquisition and analysis. We also thank Yueyi Zhang for her assistance with colony maintenance.

## Author contributions

Conceptualization, B.W. and Z.J.H.; Methodology, B.W., Z.J.H., M.S.B.S., and M.C.C. Investigation, B.W., M.S.B.S., and M.H.; Formal Analysis, B.W. and M.S.B.S. Writing – Original Draft, B.W. and Z.J.H.; Writing – Review and Editing, M.S.B.S., M.H., and M.C.C.; Funding Acquisition, Z.J.H.; Supervision, Z.J.H. and M.C.C.

## Data and materials availability

All data is available in the main text or the supplementary materials.

## Supplementary Materials

Materials and Methods

Figures S1-S12

Movies S1-S2

References (40-43)

## References

1. T. A. Seabrook, T. J. Burbridge, M. C. Crair, A. D. Huberman, Architecture, Function, and Assembly of the Mouse Visual System. Annu Rev Neurosci 40, 499–538 (2017).

2. K. S. Lee, K. Vandemark, D. Mezey, N. Shultz, D. Fitzpatrick, Functional Synaptic Architecture of Callosal Inputs in Mouse Primary Visual Cortex. Neuron 101, 421–428 e425 (2019).

3. L. Restani et al., Functional masking of deprived eye responses by callosal input during ocular dominance plasticity. Neuron 64, 707–718 (2009).

4. X. Zhao, H. Chen, X. Liu, J. Cang, Orientation-selective responses in the mouse lateral geniculate nucleus. J Neurosci 33, 12751–12763 (2013).

5. T. N. Wiesel, D. H. Hubel, Comparison of the effects of unilateral and bilateral eye closure on cortical unit responses in kittens. J Neurophysiol 28, 1029–1040 (1965).

6. T. K. Hensch, Critical period plasticity in local cortical circuits. Nat Rev Neurosci 6, 877–888 (2005).

7. J. B. Ackman, T. J. Burbridge, M. C. Crair, Retinal waves coordinate patterned activity throughout the developing visual system. Nature 490, 219–225 (2012).

8. A. Thompson, A. Gribizis, C. Chen, M. C. Crair, Activity-dependent development of visual receptive fields. Curr Opin Neurobiol 42, 136–143 (2017).

9. A. R. Woodruff, S. A. Anderson, R. Yuste, The enigmatic function of chandelier cells. Front Neurosci 4, 201 (2010).

10. H. Taniguchi, J. Lu, Z. J. Huang, The spatial and temporal origin of chandelier cells in mouse neocortex. Science 339, 70–74 (2013).

11. Q. Wang, E. Gao, A. Burkhalter, Gateways of ventral and dorsal streams in mouse visual cortex. J Neurosci 31, 1905–1918 (2011).

12. K. T. Sultan et al., Progressive divisions of multipotent neural progenitors generate late-born chandelier cells in the neocortex. Nat Commun 9, 4595 (2018).

13. F. K. Wong et al., Pyramidal cell regulation of interneuron survival sculpts cortical networks. Nature 557, 668–673 (2018).

14. D. G. Kirsch et al., p53 controls radiation-induced gastrointestinal syndrome in mice independent of apoptosis. Science 327, 593–596 (2010).

15. J. Lu et al., Selective inhibitory control of pyramidal neuron ensembles and cortical subnetworks by chandelier cells. Nat Neurosci 20, 1377–1383 (2017).

16. H. Mizuno, T. Hirano, Y. Tagawa, Evidence for activity-dependent cortical wiring: formation of interhemispheric connections in neonatal mouse visual cortex requires projection neuron activity. J Neurosci 27, 6760–6770 (2007).

17. C. L. Wang et al., Activity-dependent development of callosal projections in the somatosensory cortex. J Neurosci 27, 11334–11342 (2007).

18. H. Zhu, B. L. Roth, DREADD: a chemogenetic GPCR signaling platform. Int J Neuropsychopharmacol 18, (2014).

19. J. Olavarria, R. Malach, R. C. Van Sluyters, Development of visual callosal connections in neonatally enucleated rats. J Comp Neurol 260, 321–348 (1987).

20. L. A. Kirkby, G. S. Sack, A. Firl, M. B. Feller, A role for correlated spontaneous activity in the assembly of neural circuits. Neuron 80, 1129–1144 (2013).

21. D. A. Arroyo, M. B. Feller, Spatiotemporal Features of Retinal Waves Instruct the Wiring of the Visual Circuitry. Front Neural Circuits 10, 54 (2016).

22. F. M. Rossi et al., Requirement of the nicotinic acetylcholine receptor beta 2 subunit for the anatomical and functional development of the visual system. Proc Natl Acad Sci U S A 98, 6453–6458 (2001).

23. A. Tiriac, B. E. Smith, M. B. Feller, Light Prior to Eye Opening Promotes Retinal Waves and Eye-Specific Segregation. Neuron 100, 1059–1065 e1054 (2018).

24. J. A. Gordon, M. P. Stryker, Experience-dependent plasticity of binocular responses in the primary visual cortex of the mouse. J Neurosci 16, 3274–3286 (1996).

25. B. S. Wang, R. Sarnaik, J. Cang, Critical period plasticity matches binocular orientation preference in the visual cortex. Neuron 65, 246–256 (2010).

26. L. Baroncelli, C. Braschi, L. Maffei, Visual depth perception in normal and deprived rats: effects of environmental enrichment. Neuroscience 236, 313–319 (2013).

27. J. Faguet, B. Maranhao, S. L. Smith, J. T. Trachtenberg, Ipsilateral eye cortical maps are uniquely sensitive to binocular plasticity. J Neurophysiol 101, 855–861 (2009).

28. J. T. Trachtenberg, Competition, inhibition, and critical periods of cortical plasticity. Curr Opin Neurobiol 35, 44–48 (2015).

29. A. K. Engel, P. Konig, A. K. Kreiter, W. Singer, Interhemispheric synchronization of oscillatory neuronal responses in cat visual cortex. Science 252, 1177–1179 (1991).

30. D. C. Kiper, M. G. Knyazeva, L. Tettoni, G. M. Innocenti, Visual stimulus-dependent changes in interhemispheric EEG coherence in ferrets. J Neurophysiol 82, 3082–3094 (1999).

31. L. G. Nowak, M. H. Munk, J. I. Nelson, A. C. James, J. Bullier, Structural basis of cortical synchronization. I. Three types of interhemispheric coupling. J Neurophysiol 74, 2379–2400 (1995).

32. J. C. Gardner, M. S. Cynader, Mechanisms for binocular depth sensitivity along the vertical meridian of the visual field. Brain Res 413, 60–74 (1987).

33. C. Peiker, T. Wunderle, D. Eriksson, A. Schmidt, K. E. Schmidt, An updated midline rule: visual callosal connections anticipate shape and motion in ongoing activity across the hemispheres. J Neurosci 33, 18036–18046 (2013).

34. N. L. Rochefort et al., Functional selectivity of interhemispheric connections in cat visual cortex. Cereb Cortex 19, 2451–2465 (2009).

35. K. E. Schmidt, D. S. Kim, W. Singer, T. Bonhoeffer, S. Lowel, Functional specificity of long-range intrinsic and interhemispheric connections in the visual cortex of strabismic cats. J Neurosci 17, 5480–5492 (1997).

36. N. Dehorter, L. Vinay, C. Hammond, Y. Ben-Ari, Timing of developmental sequences in different brain structures: physiological and pathological implications. Eur J Neurosci 35, 1846–1856 (2012).

37. J. F. Sauer, M. Bartos, Postnatal differentiation of cortical interneuron signalling. Eur J Neurosci 34, 1687–1696 (2011).

38. C. Rock, A. J. Apicella, Callosal projections drive neuronal-specific responses in the mouse auditory cortex. J Neurosci 35, 6703–6713 (2015).

39. M. G. Shuler, D. J. Krupa, M. A. Nicolelis, Bilateral integration of whisker information in the primary somatosensory cortex of rats. J Neurosci 21, 5251–5261 (2001).

40. J. B. Ackman, H. Zeng, M. C. Crair, Structured dynamics of neural activity across developing neocortex. bioRxiv, 012237 (2014).

41. X. An, H. Gong, N. McLoughlin, Y. Yang, W. Wang, The Mechanism for Processing Random-Dot Motion at Various Speeds in Early Visual Cortices. PLOS ONE. 9, e93115 (2014).

42. C. A. Leamey et al., Ten_m3 regulates eye-specific patterning in the mammalian visual pathway and is required for binocular vision. PLoS Biol. 5, e241 (2007).

43. A. D. Huberman, M. B. Feller, B. Chapman, Mechanisms underlying development of visual maps and receptive fields. Annual review of neuroscience. 31, 479–509 (2008).

