## Supplementary Materials for "Retinal and callosal activity-dependent chandelier cell elimination shapes binocularity in primary visual cortex"

**This PDF file includes:**

Materials and Methods

Figs. S1 to S12

Captions for Movies S1 to S2

**Other Supplementary Materials for this manuscript include the following:**

Movies S1 to S2

### Materials and Methods

Experimental animals

In order to genetically label and manipulate chandelier cells, we crossed *Nkx2.1 -CreER* mice (The Jackson Laboratory stock 014552) with either *Rosa26-lox-stop-lox-TdTomato (Ai14)* reporter (The Jackson Laboratory stock 007914). In order to properly identify embryonic day 17.5 (E17.5) for tamoxifen (TM) inductions, Swiss Webster or C57B6 (Taconic) females were housed with *Nkx2.1CreER;Ai14 (het/homo)* males overnight and females were checked for vaginal plug by 8-9 am the following morning. Positive plug identification was timed at E0.5. Emx1-cre/Ai95 mice were used for *in vivo* calcium imaging (Emx1-cre, The Jackson Laboratory 005628; Ai95, provided by Hongkui Zeng and available as Ai95D from The Jackson Laboratory 028865). The in-vivo calcium imaging experiments were performed in compliance with the Yale IACUC, U. S. Department of Health and Human Services and Institution guidelines. The rest of the experiments were conducted in accordance with the Institutional Animals Care and Use Committee of Cold Spring Harbor Laboratory.

TM induction TM was dissolved in corn oil (20 mg/ml) overnight, at room temperature under constant stirring. Stocks were stored as individual aliquots at 4°C degrees for no more than one month. Following light isoflurane anesthesia pregnant females were given oral gavage administration of TM (dose: 3 mg / 30 g of body weight) at gestational day E17.5 (embryonic day 17.5) for dense revealing of ChCs. In rare instances, TM induction lead to dystocia in pregnant females and emergency caesarian sections were performed. Pups retrieved following caesarians were housed with Swiss Webster foster mothers until weaning age.

Viral Constructs

*AAV-CB7-EGFP* was purchased from Penn Vector Core, Philadelphia, PA. *AAV-Hsyn-HA-HM4D(Gi)-IRES-mCitrine* was purchased from UNC vector core, Chapel Hill, North Carolina. *AAV-hSyn1-HA-hM4D(Gi)-IRES-mCitrine-WPRE-hGHp(A)* from ETH Zurich stock # V 92-8. *AAV8-hSyn-DIO-hM4D(Gi)- mCherry* (Addgene, plasmid #44362, lots v9165 and v24795).

DNA Plasmids

CAG-Kir2.1 plasmid was a gift from Yu-Qiang Ding. pCAGGS-IRES-EGFP plasmid was a gift from Linda Van Aelst. pAAV-hSyn-HA-hM4D(Gi)-IRES-mCitrine was a gift from Bryan Roth (Addgene plasmid # 50464 ; http://n2t.net/addgene:50464 ; RRID:Addgene_50464). pAAV-CAGGS-HA-hM4D(Gi)-IRES-mCitrine plasmid cassettes was assembled and cloned using standard molecular cloning protocols with restriction enzymes from New England Biolabs. CMV-CAAGS promoter was subcloned into pAAV-Hsyn-HA-hM4D(Gi)-IRES_mCitrine replacing Hsyn promoter. All constructs were sequenced to ensure their fidelity and proper reversed orientation of given inserts

In Utero Electroporation

Timed pregnant female mice (crossed to were *Nkx2.1 –CreER*;Ai14) were used for in utero electroporation. The animal was anesthetized, the abdominal cavity was cut open, and the uterine horns were exposed. Approximately 1–2 µg DNA was injected into the lateral ventricle at E15–E16 by using a pulled glass micropipette. The angle of electrical paddles was adjusted to target visual cortex. Square electric pulses (45 V, 50 ms) were passed five times at 1-s intervals.

Embryos were put back into the cavity as soon as the electroporation was done. Embryos were allowed to develop normally to birth.

Surgical Procedures of stereotaxic injection: Adult

Adult animals were anesthetized by inhalation of isoflurane (2%). Mice were mounted in a stereotaxic headframe (Kopf Instruments Model 940 series). Coordinates were identified for primary visual cortex (Adults: 3.0–3.3 mm lateral from the midline and 0.5–0.8 mm anterior from the lambda suture). An incision was made over the scalp, a small burr hole was made into the skull and brain surface was exposed. A pulled glass pipette tip of 20-30 μm containing virus was lowered into the brain. Pulses were delivered using a Picospritzer (General Valve Corp) at a rate of 30 nl/minute; the pipette was left in the brain for 5-10 minutes to prevent backflow. Following the injection, the pipette was withdrawn, the incision was closed with tissue glue, and animals recovered.

Surgical Procedures of stereotaxic injection: P0-P1 pups

P0-1 mice underwent hypothermia anesthesia in preparation for surgery. They were mounted and stabilized in a neonatal injection setup developed in-house. Neonates have closed ear canals and earbars cannot be used for head stabilization and alignment. A small scalp incision was made and V1 was targeted with a MP285 micromanipulator (Sutter Instruments). The targeted coordinates are 1.5mm lateral from the midline and 0.5mm anterior from the lambda suture. A pulled glass capillary (3.5” #3-000-203-G/X capillary, Drummond Scientific Co, or G150TF-4 capillary, Warner Instruments; P-97 pipette puller, Sutter Instruments) mounted on a Nanjoject III (Drummond Scientific Co) or a Picospritzer (General Valve Corp) was filled with virus and slowly lowered until piercing the surface of the skull. The capillary was raised to the desired depth, followed by a waiting period of 30-60 seconds, and 200-250nl were injected at rate of 1-2nl/s. Once the injection was completed, there was a 30-60s waiting period followed by capillary withdrawal and closing of the incision with tissue glue (Vetbond). Pups recovered on a heating pad before returning to the home cage.

Surgical procedure for *in vivo* imaging

P9-P14 mice were deeply anesthetized with 2% isoflurane, received local analgesia (0.5% lidocaine) and maintained on a heated water pad (HTP-1500, Adroit Medical Systems). Scalp and connective tissue were removed to expose the skull, headbars placed over the occipital and nasal bones, and cyanoacrylate glue applied to headfix the animal and cover the skull. Imaging began after 3h+ of recovery from anesthesia and the animal was periodically hydrated with subcutaneous saline. After 1h+ of spontaneous activity recordings, control animals received a single CNO (10mg/Kg) i.p. injection to assess acute effects.

Histology

For histology, animals were perfused with 4% PFA in PBS. Brains were removed and postfixed overnight in same fixative. Coronal brain slices were sectioned at a 75 µm thickness via vibratome. Sections were blocked with 10% normal goat serum in 0.5% Triton in PBS and then incubated overnight with combinations of the following primary antibodies diluted in block solution: rabbit polyclonal RFP (1:1000, Rockland) or chicken polyclonal anti-GFP (1:1000, Aves) for fluorophore preservation of tdTomato (from *Ai14*) and GFP virus expression, mouse monoclonal anti-parvalbumin (1:1000 Sigma), rabbit polyclonal Calretinin (1:250, Immunostar), rat monoclonal anti-muscarinic acetylcholine receptor m2 (m2AchR) (1:500, Millipore Sigma), and rabbit 5-hydroxytryptamine (5-HT) (1:1000 Immunostar). Sections were incubated with appropriate Alexa fluor dye-conjugated IgG secondary antibodies (1:500, Molecular Probes). Sections were washed and mounted with Fluoromount-G (Southern Biotech).

Image Acquisition and Analysis

Images were taken by confocal microscopy (Zeiss LSM 780). All images were processed using Fiji. To determine the callosal projecting region, Z-stack images (10μm optical sections for 75 μm sections) were acquired with a 10X objective. Maximum intensity projections of the Z stacks were obtained. Region of callosal axon projections at V1/V2L border was selected and threshold at 30% of peak intensity amplitude to determine the area of callosal projection zone. In the Kir2.1 treated animals were the callosal projections were disrupted, we used the expression of m2AchR to determine the callosal projection zone. First we identify the lateral most callosal projection zone at V1/V2L border in control animals and correlate that point with the m2AchR expression profile. On average, the m2AchR expression reduces to 93% of the peak intensity at the lateral edge of callosal projection zone. The average width of callosal projection zone is 272μm ± 5μm. Then, in Kir2.1 treated animals we determine when m2AchR expression reduces to 93% of the peak intensity as the lateral edge of callosal projection zone and used 272μm ± 5μm as the width of the zone. For estimating callosal projection zone at P7 when callosal axons have not reached contralateral cortex, we used the expression of 5-hydroxytryptamine (5-HT) to identify V1 and estimate the lateral most 200μm as the V1/V2L border that receive callosal input.

Intravitreous injection

Every 24 hr from P7 to P14, mice were anesthetized using isoflurane (2%), and 1 μl of 10 uM tetrodotoxin (Sigma, MO) in sterile $H_2O$, or sterile $H_2O$ alone, was injected intravitreously into one ore both eyes. Eye injections at P5 and P7 the eyelids were cut open. A 34G needle attached to a Hamilton (Reno, NV) microsyringe was used to inject the solution at the rate of about 1 μl/min into the vitreous humor at the ora serrata. The needle is withdrawn after holding it in place for 30 s to 1 min. The animals were allowed to survive to P40–P60 before analysis.

Monocular Enucleation

For monocular enucleation in the first few days postnatally (P0-P1), we will anesthesia the pups through cooling; otherwise, for monocular enucleationat P6, the animal is asnestheized using isoflurane (2%). The eyelids will be gently pried open (the eyes open at day 13) to expose the globe. Curved forceps will be used to gain access to the perimeter and back of the globe. The globe is removed whole after the posterior bundle containing the optic nerve and artery are pinched and transected. The eyelids are then repositioned and sealed using ophthalmic surgical adhesive. After recovery from anesthesia, the animals will be returned to their mothers. This surgery requires less than one minute.

Chemical-genetic manipulation

For the chemical-genetic manipulation, *NkxCre-ER;Ai14* mice that received injections of either the AAV/plasmid form of hM4Di-mCitrine or the EGFP (control) into visual cortex were intraperitoneally (i.p.) injected with CNO (10 mg/kg) or saline.

Widefield Calcium imaging and assessment of mCherry-expression

Widefield calcium imaging was performed using a Zeiss Axiozoom V.16 with PlanNeoFluar Z 1X, 0.25 NA objective and equipped with an ET-EGFP filter (Chroma, 49002). Epifluorescence excitation was delivered with a 460nm LED source (X-cite XLED1, Excelitas Technologies). Emissions were collected with a sCMOS camera (pco.edge 4.2) with 512x500 resolution after 4x4 pixel binning, and frames acquired using Camware software (pco). Each recording lasted 10 minutes at 10 frames/second (100ms exposure time), and at least 60 minutes of data were collected per animal. *AAV8-hSyn-DIO-hM4D(Gi)- mCherry* was confirmed using the same setup for widefield calcium imaging, a 565nm LED source (X-cite XLED1, Excelitas Technologies) and an ET-AlexFluor568 filter set (Chroma 49031).

Calcium signal detection and analysis

Image processing and calcium signal detection was performed using the freely available wholeBrainDX software developed by James Ackman (available at http://github.com/ackman678/wholeBrainDX) and written for MATLAB (MathWorks) (40) (Gaussian filter(σ) = 3px, dF/F threshold = 2 SD). Visual Cortex ROIs were individually defined using functional data (domain activation frequency and/or duration maps), hand-drawn on ImageJ software and divided into lateral and medial regions according to V1's apex and base. Activations were assigned to ROIs based on their centroid, and activation frequency was calculated as the mean number of individual activations per minute, averaged across recordings.

We performed a variety of seed-based correlations to quantitatively measure the spatial extent of similar V1 activity within- and across- hemispheres. Seed-based correlation analyses reveal regions with similar activity patterns, where tight correlations are interpreted as representing similar functional roles. Individual seed-based correlation maps were computed for seeds from a grid covering lateral V1 of the DREADD-injected hemisphere (range = 29-51 seeds, mean=36.52, median=35, fig. S6F) Maps from individual recordings were averaged to obtain a mean correlation map per seed. For each mean map, we calculated the proportion of each V1 region tightly correlated with the seed (pearson's correlation coefficient ≥ 0.7), then averaged these values across maps, and normalized them relative to the seed region (V1 lateral of the DREADD-injected hemisphere). All averaging and normalizations were done within animal.

In vitro Electrophysiology

*Slice preparation.* We used wildtype mice electroporated with pAA-CAGGS-HA-hM4D(Gi)-IRES-mCitrine at E15.5 to confirm suppression of neuronal activity by chemical genetic method. Mice (~P7) were anesthetized with isoflurane before decapitation.

The dissected brain was rapidly immersed in ice-cold, oxygenated, artificial cerebrospinal fluid (section ACSF: 110 mM choline-Cl, 2.5 mM KCl, 4mM MgSO4, 1mM CaCl2, 1.25 mM NaH2PO4, 26mM NaHCO3, 11mM D-glucose, 10 mM Na ascorbate, 3.1 Na pyruvate, pH 7.35, 300 mOsm) for 1 min. Coronal prefrontal cortical slices were sectioned at 300 μm thickness using a vibratome (HM 650 V; Microm) at 1-2 °C and incubated with oxygenated ACSF (working ACSF; 124mM NaCl, 2.5 mM KCl, 2 mM MgSO4, 2 mM CaCl2, 1.25 mM NaH2PO4, 26 mM NaHCO3, 11 mM D-glucose, pH 7.35, 300mOsm) at 34 °C for 30 min, and then transferred to ACSF at room temperature (25 °C) for >30 min before use. Whole cell patch recordings were directed electroporated hemisphere, the morphology of subcortical whiter matter and corpus callosum as primary landmarks according to the atlas (Paxinos and Watson Mouse Brain in Stereotaxic Coordinates, 3rd edition).

Patch pipettes were pulled from borosilicate glass capillaries with filament (1.2 mm outer diameter and 0.69 inner diameter; Warner Instruments) with a resistance of 3-6 MΩ. The pipette recording solution consisted of 130 mM potassium gluconate, 15 mM KCl, 10 mM sodium phosphocreatine, 10 mM Hepes, 4 mM ATP·Mg, 0.3 mM GTP, and 0.3 mM EGTA (pH 7.3 adjusted with KOH, 300 mOsm). Whole cell recordings from GFP labeled cells in layer 2/3 were made with Axopatch 700B amplifiers (Molecular Devices, Union City, CA) using an upright microscope (Olympus, Bx51) equipped with infrared-differential interference contrast optics (IR-DIC) and fluorescence excitation source. GFP negative cells were blindly selected within 100 um distance to GFP positive cells as controls. Both IR-DIC and fluorescence images were captured with a digital camera (Microfire, Optronics, CA). All recordings were performed at 33–34 °C with the chamber perfused with oxygenated working ACSF. Recordings were made with two MultiClamp 700B amplifiers (Molecular Devices). The membrane potential was maintained at -75mV in the voltage clamping mode and zero holding current in the current clamping mode, without the correction of junction potential. Signals were recorded and filtered at 2 kHz, digitalized at 20 kHz (DIGIDATA 1322A, Molecular Devices) and further analyzed using the pClamp 10.3 software (Molecular Devices) for intrinsic properties and synaptic features.

In vivo Electrophysiology

*Nkx2.1CreER;Ai14* mice (TM induction at E17.5) age between 2-4 months were placed in a stereotaxic apparatus on a heating pad. The animal's temperature was monitored and maintained at 37°C through a feed-back heater control module (Frederick Haer Company, Bowdoinham, ME). Silicon oil was applied to both eyes to prevent them from drying. The animals were lightly anesthetized under isoflurane anesthesia. A small craniotomy (2 $mm^2$) was performed at the left hemisphere to expose the cortex for recording. A 32-Channel electrode array consisting of 8 tetrodes (A4x2-tet-5mm-150-200-121-A32, Neuronexus) was penetrated perpendicular to the pial surface in the binocular zone of V1 (3.0-3.3 mm lateral from the midline and 0.5-0.8 mm anterior from the lambda suture). Only cells in upper layer 2-3 were recorded and analyzed. Action potentials were recorded extracellularly (sampled at 32 kHz) with Cheetah32 system (Neuralynx, Inc.). The animals were euthanized at the end of the recording.

All data analysis was carried out using built-in and custom-built software in Matlab (Mathworks). Spikes were manually sorted into clusters (presumptive neurons) off-line based on peak amplitude and waveform energy using the MClust software (A.D. Redish). Cluster quality was quantified using isolation distance and L-ratio. Putative cells with isolation distance < 20 or L-ratio > 0.1 were excluded.

Visual stimuli was generated with Matlab programs developed by Dr. Xu An using the Psychophysics Toolbox extensions (41). An Asus monitor (120 Hz refresh rate) was placed at 25 cm in front of the animal to display the stimuli, with its midline aligned with the animal. Stimuli were presented to either eye separately with the other eye occluded. Sinusoidal grating drifting perpendicular to their orientation were used to determine V1 neuron's orientation selectivity and spatial tuning. The drifting direction and spatial frequency of the gratings (full contrast and temporal frequency of 2 Hz) were varied between $0^o$ – $330^o$ (12 steps at $30^o$ spacing) and 0.01-0.08 cycle/degree (in 4 logarithmic steps) in a pseudorandom order. The mean firing rate during the period of stimulus presentation was used to generate the direction tuning curves. The preferred direction was determined as the one that gave maximum response ($R_{pref}$), averaged across all spatial frequencies. The preferred spatial frequency (pref_SF) was the one that gave peak response at this direction. A vector summation method was used to quantitatively characterize the direction tuning curves:

$$S = \frac{\sum_k r_K e^{i\theta_k}}{\sum_k r_K},$$

Where $\theta_k$ and $r_K$ are the direction of motion and mean firing rate, respectively. The complex phase and amplitude of the resultant S represent the preferred direction (pref-D) and direction selective index (DSI), respectively. DSI varies between 0, for a cell that responds equally to all directions, and 1, for a cells that only responds to a single direction. Preferred orientation (pref_O) and orientation selective index (OSI) to sine-wave grating were calculated by substituting $2\theta_k$ for $\theta_k$, while $\pi$ was added to the resultant phase before it was halved as the moving directions of grating were always perpendicular to the orientations. The difference in preferred orientation between the two eyes was calculated by subtracting ipsilateral pref_O from contralateral pref_O along the 180° cycle (-90° to 90° ). The absolute values of these differences ($\Delta O$) were used in all quantifications for binocular matching of orientation preference. The ODI for each cell was calculated as $(C - I)/(C + I)$, where $C$ and $I$ represent the maximum response magnitude for the contralateral and ipsilateral eyes, respectively. The ODI ranges from -1 to 1, where positive values indicate contralateral bias and negative values indicate ipsilateral bias.

Visual Cliff Test

For the visual cliff test, mice were placed in a 61 cm X 61 cm box with a clear acrylic (ShopPoPdisplays) base. A high-contrast grating was attached to the underside of one half of the box. The box placed such that the clear half of the base protruded from a table, revealing a drop to the floor of approximately 100 cm. Mice were placed in the center of the box and their behavior monitored for 10 min via a digital video camera mounted above the floor of the box. The data were analyzed, blind to genotype, from video recordings. An approach was defined as moving from the patterned region towards the ‘‘cliff’’ with the nose of the animal within 5 cm of the midline. Crossing was defined as the animal completely crossing the midline from the patterned to the clear half of the box. Retreating was defined as moving such that the head was within 5 cm of the midline and retreating without the entire body crossing the border (42). Subsequent approaches were not scored until the mouse had moved outside the 5-cm range. The analysis was further divided into middle crossing and side crossing, where the middle crossing was scored when the animal was more than 5cm away from the sidewalls while the side crossing was scored when the animal crossed the cliff within 5 cm from the sidewalls.

Analysis and Statistics

In-vivo electrophysiology and calcium imaging summary data values were obtained with MATLAB 2017a (MathWorks). All data were tested and plotted using GraphPad Prism 8.0.2 for Windows (GraphPad Software) or MATLAB 2017a (MathWorks). For comparison of two groups of data, KS tests, unpaired, and paired Student's t tests were used as indicated in the experiment. Data are presented as mean ± s.e.m or mean ± SD (for in-vivo calcium imaging data). For in-vivo calcium imaging data, statistical tests were 2-way ANOVAS followed by Sidak's multiple comparisons tests. If not specifically indicated, and a p value $<0.05$ was considered significant. The significance was marked as *, $p < 0.05$; **, $p <0.01$ and ***, $p < 0.001$.

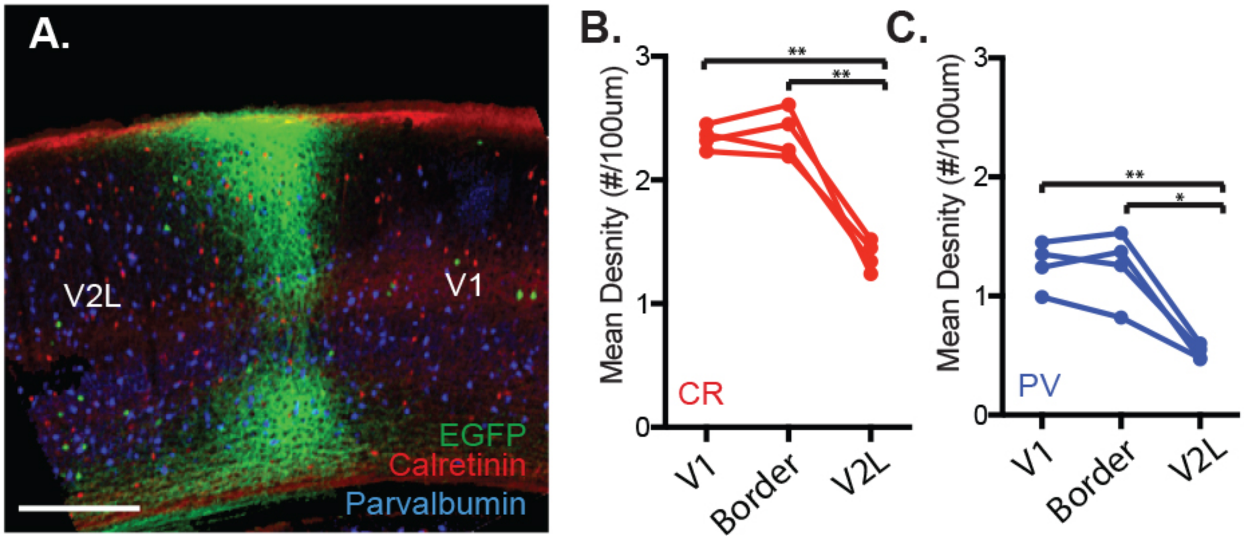


**Fig. S1. Parvalbumin or Calretinin positive cells are equally distributed across V1 and V1/V2L border**

(**A**) Distribution of parvalbumin (PV) or calretinin (CR) positive cells at the V1/V2L border. The border is identified by AAV-EGFP tagged callosal projection axons. (**B-C**) Mean CR (**B**) or PV (**C**) cell density to V1 showing similar density of these cells between V1/V2L border and V1 (n = 4 animals). * $p<0.05$, ** $p<0.01$. All statistics performed with paired t-test unless stated otherwise. Scale Bar: 200um.

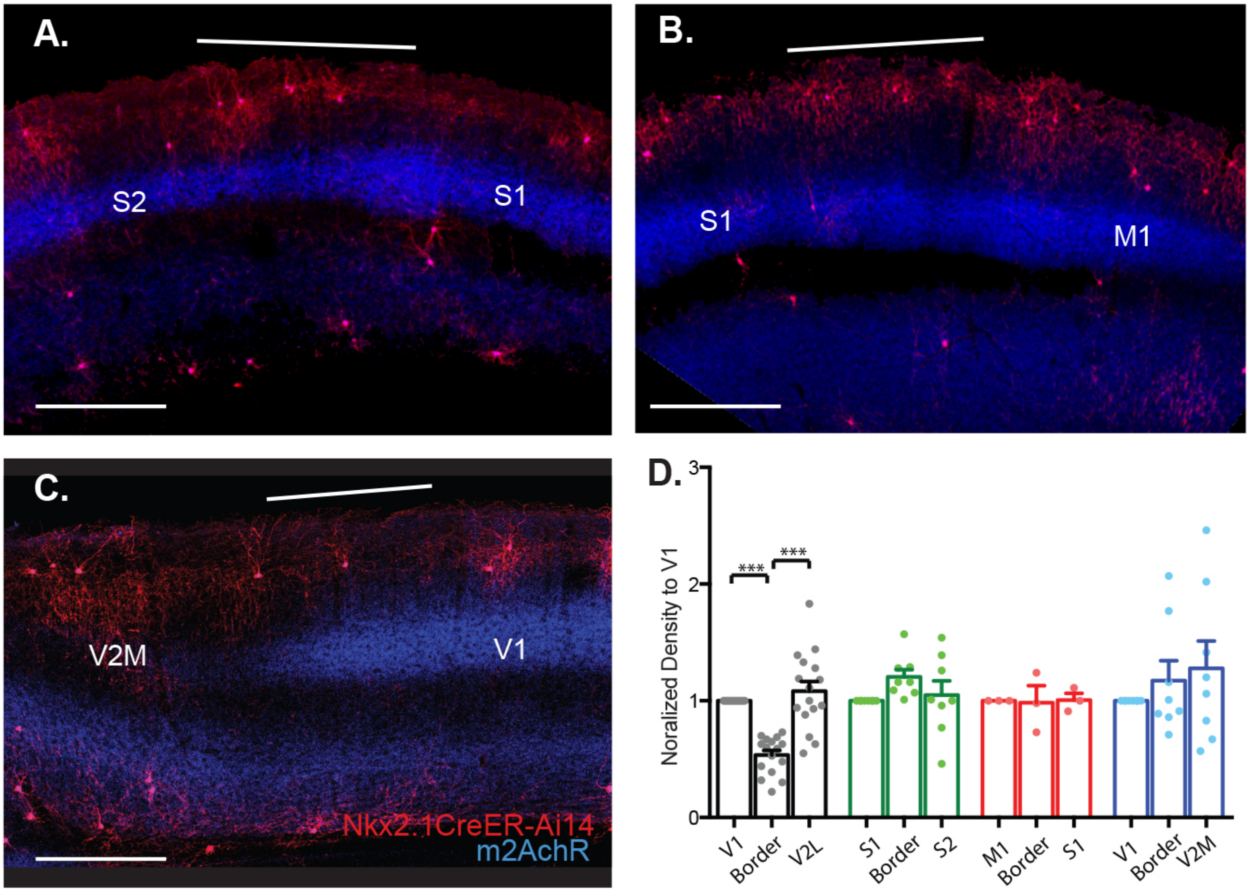


**Fig. S2. Chandelier cell distribution is similar across other border regions.**

(**A**) Example image of Chandelier cell (ChC) distribution across primary (S1) and secondary (S2) somatosensory cortex. (**B**) Example image of ChC distribution across S1 and primary motor (M1) cortex. (**C**) Example image of ChC distribution across V1 and medial secondary visual cortex (V2M). (**D**) Normalized ChC density to primary cortical area showing similar ChC distribution across border regions between S1-S2 (n = 8 animals), M1-S1 (n = 3 animals), and V1-V2M (n = 8 animals). * $p < 0.05$, ** $p < 0.01$, *** $p < 0.001$. All statistics performed with paired t-test unless stated otherwise. Scale Bar: 200um.

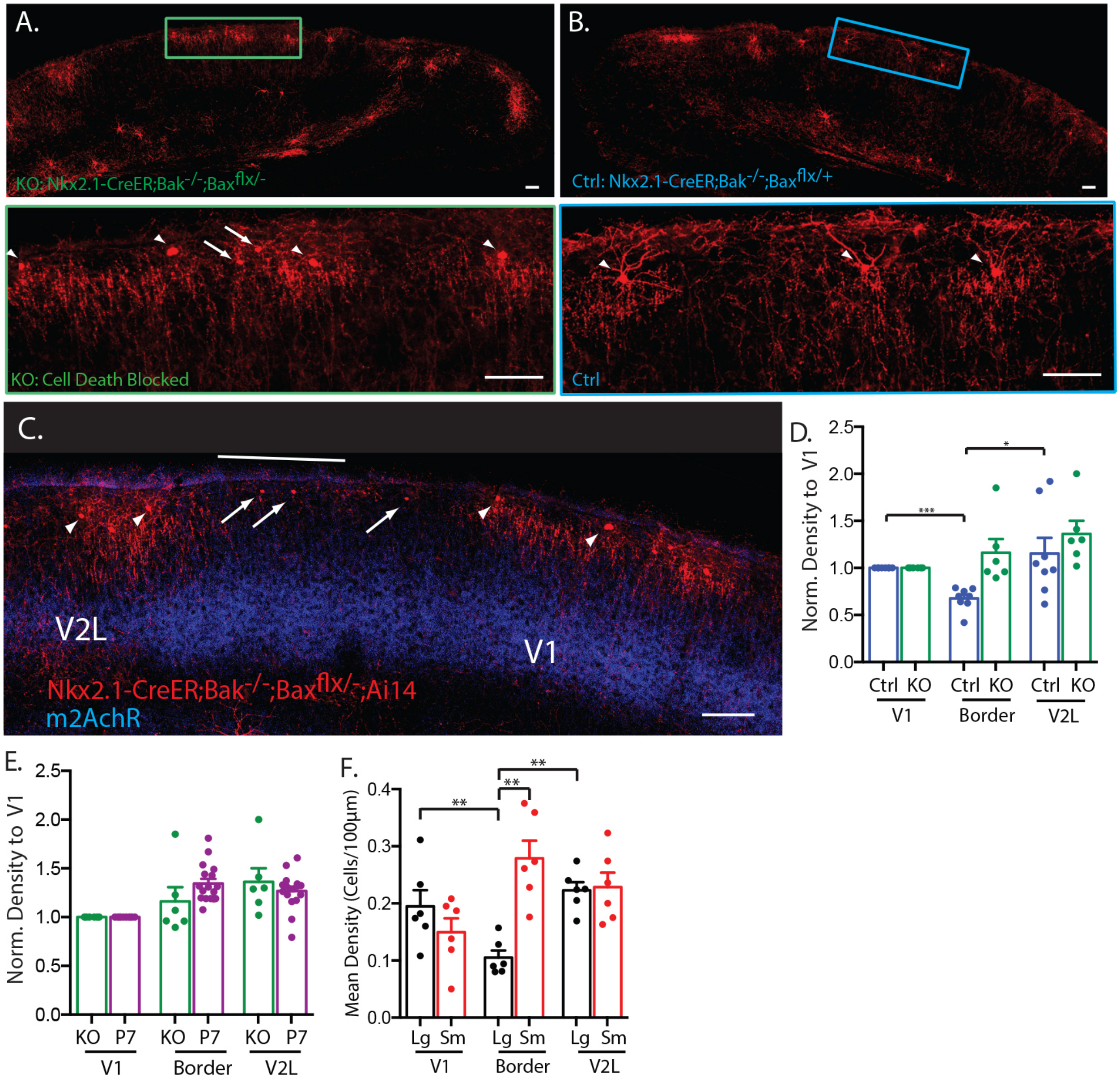


**Fig. S3. Developmental Chandelier cell elimination is mediated through apoptosis**

(**A-B**) Blockade of apoptosis in visual cortex of Nkx2.1-CreER;Bak$^{-/-}$;Bax$^{f/-}$ mouse resulted in more ChCs (**A**) than controls (**B**). (**C**) More ChCs remain at V1/V2L border in apoptosis blocked mice. (**D**) More ChCs remain at V1/V2L border in KO animals (n = 6 animals) than the controls (n = 8 animals). (**E**) The distribution pattern of ChCs in KO adult animals closely resembles the pattern in wild type P7 mice before ChC elimination. (**F**) Most of the remaining RFP+ cells at the border are small atrophic ChCs that are destined to die but are blocked from apoptosis. Arrows point to small atrophic ChCs (sm). Arrowheads point to normal ChCs (Lg). * $p < 0.05$, ** $p < 0.01$, *** $p < 0.001$. All statistics performed with paired t-test unless stated otherwise. Scale Bar: 100um.

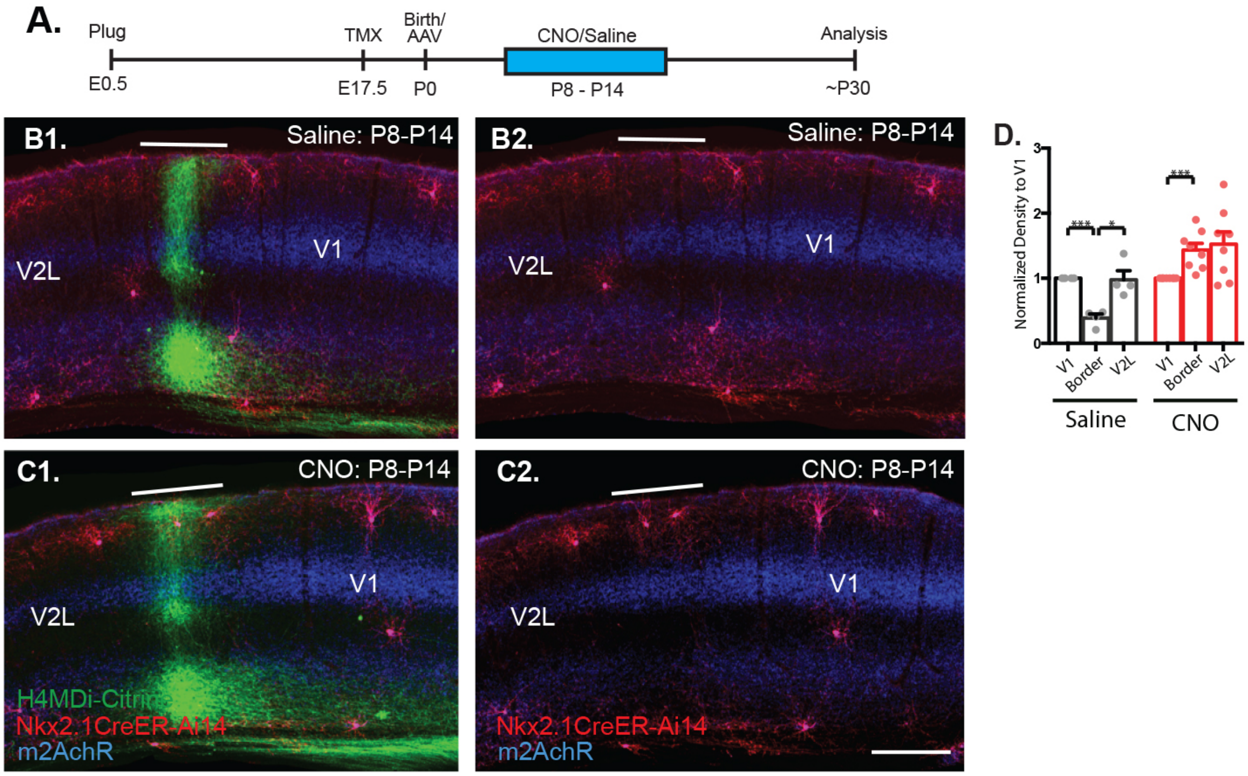


**Fig. S4. AAV injection of Hm4Di at P0 and CNO application between P8-P14 prevents Chandelier cell elimination at V1/V2L border.**

(**A**) Time course of tamoxifen induction at E17.5, AAV-Hm4Di-mCitrine injection at P0, and daily CNO or saline injection between P8-P14. (**B-C**) Example image of ChC distribution at the border region between V1 and V2L in saline treated control (**B**) and CNO treated (**C**) animals. (**D**) Decrease in ChC density at V1/V2L border in saline control (n = 4 animals), but not binocular CNO treated animals (n = 8 animals). * $p < 0.05$, ** $p < 0.01$, *** $p < 0.001$. All statistics performed with paired t-test unless stated otherwise. Scale Bar: 200um.

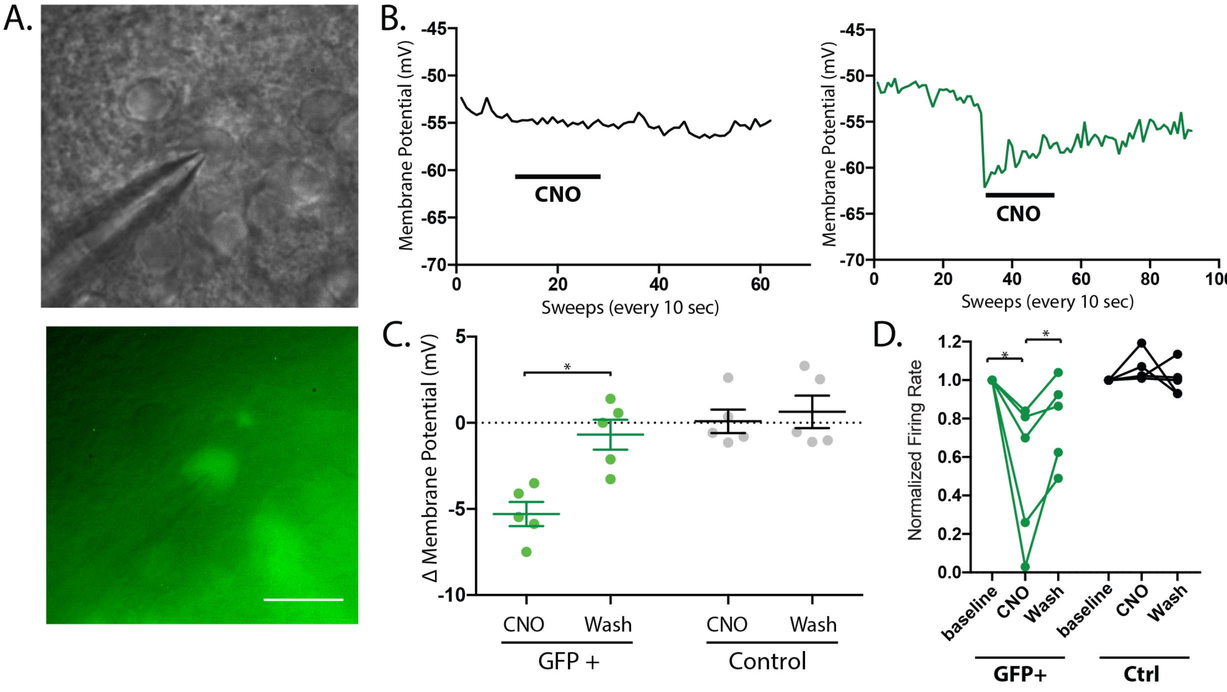


**Fig. S5. Suppression of neuronal activity by chemical genetic method.**

(**A**) An example of whole-cell patchgin recording of a hM4Di-mCitrine labeled cell. (**B**) A representative trace recording in current clamp mode from a GFP-negative cell (black trace, left), or a GFP-positive cell that expresses hM4Di (green trace, right). Application of CNO (bath application for 4 mins as indicated by the bar) induced hyperpolarization in GFP-positive cell but not in GFP-negative cell. (**C**) On average, CNO induced an average of 5.3±1.6 mV (n = 5 cells) hyperpolarization that was reversible upon CNO withdrawal, whereas CNO does not induce hyperpolarization in GFP-negative controls (mean Δ membrane potential: 0.1±1.5 mV, n = 5 cells). (**D**) hM4Di activation suppresses neuronal firing in GFP+ cells but not GFP- cells. Firing rate was normalized to baseline (n = 5 cells, *$P < 0.05$). All statistics performed with paired t-test unless stated otherwise. Scale Bar: 25um.

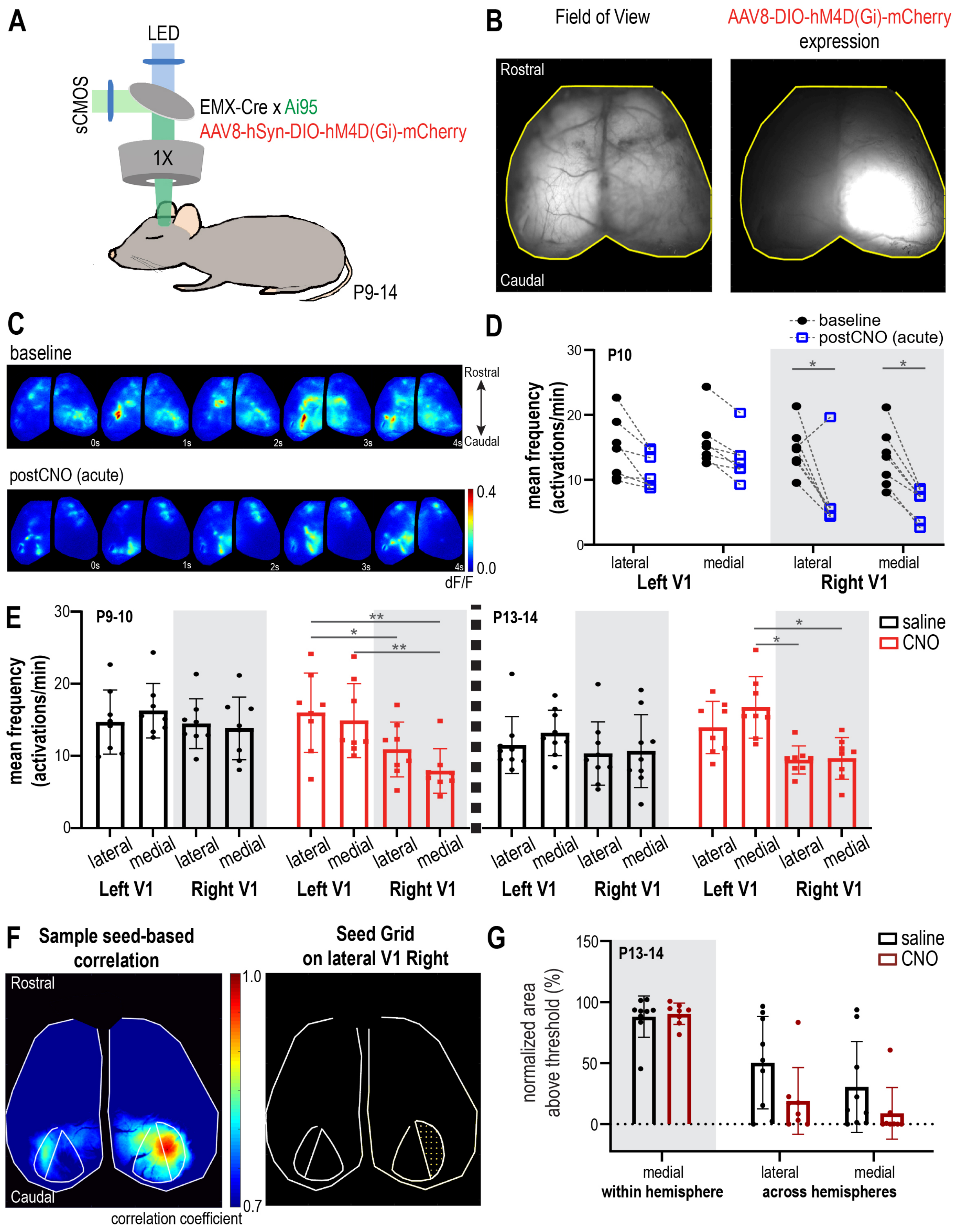
A
LED
sCMOS
EMX-Cre x Ai95
AAV8-hSyn-DIO-hM4D(Gi)-mCherry
1X
P9-14
B
Field of View
AAV8-DIO-hM4D(Gi)-mCherry expression
Rostral
Caudal
C
baseline
postCNO (acute)
0s
1s
2s
3s
4s
Rostral
Caudal
0.4
0.0
dF/F
D
baseline
postCNO (acute)
P10
mean frequency (activations/min)
lateral
medial
lateral
medial
Left V1
Right V1
E
P9-10
P13-14
saline
CNO
mean frequency (activations/min)
lateral
medial
Left V1
Right V1
F
Sample seed-based correlation
Seed Grid on lateral V1 Right
Rostral
Caudal
1.0
0.7
correlation coefficient
G
P13-14
saline
CNO
normalized area above threshold (%)
medial
within hemisphere
lateral
medial
across hemispheres

**Fig. S6. Acute and chronic CNO change cortical activity *in-vivo***

**(A)** Diagram of 1-photon mesoscale calcium-imaging set-up used for data collection (between P9-14) **(B) Left**, example field of view (FOV) of the dorsal surface of the mouse brain. **Right**, same FOV showing AAV8-DIO-hM4D(Gi)-mCherry expression (P0-1 cortical injection). **(D)** Mean activation frequency before (black) and after (blue) acute CNO administration, per cortical region (Left and Right V1, lateral and medial V1) (P10, n=7). Shaded area corresponds to data from the DREADD-injected hemisphere. Lateral V1 encompasses the binocular zone. **(E)** Effects of chronic saline (black) or CNO (red) on mean activation frequency at P9-10 (left, saline n=8, CNO n=8) or P13-14 (right, saline n=9, CNO n=8). Data is plotted per cortical region (Left and Right V1, lateral and medial V1). Shaded area corresponds to data from the DREADD-injected hemisphere and dashed column divides age groups. Please note P9-10 saline data includes the baseline data from **D**.**(F) Left**, sample seed-based correlation map for a single seed on lateral V1 Right (V1 outlined in white). **Right**, Sample grid of all seeds on lateral V1 Right for one animal. Correlation maps were computed for each seed, and similar grids and correlation map were created for every animal. **(G)** The mean proportion of each visual region highly correlated with lateral V1 Right, normalized by lateral V1 Right (DREADD-injected hemisphere, correlation coefficient threshold = 0.7, P13-14). Data is plotted for saline (black, n=9) and CNO (burgundy, n =8) as within- (shaded region) or across hemispheres. A 2-way ANOVA revealed a significant interaction between treatment and visual region ($F = 4.44$, $p = 0.021$), consistent with reduced inter-hemispheric activity correlations without effecting within hemisphere correlations. All barplots represent the mean ± SD, while circles represent individual animals, and all statistical analyses were 2-way ANOVAs with Sidak corrections for multiple comparisons, * = $p < 0.05$, ** = $p < 0.01$).

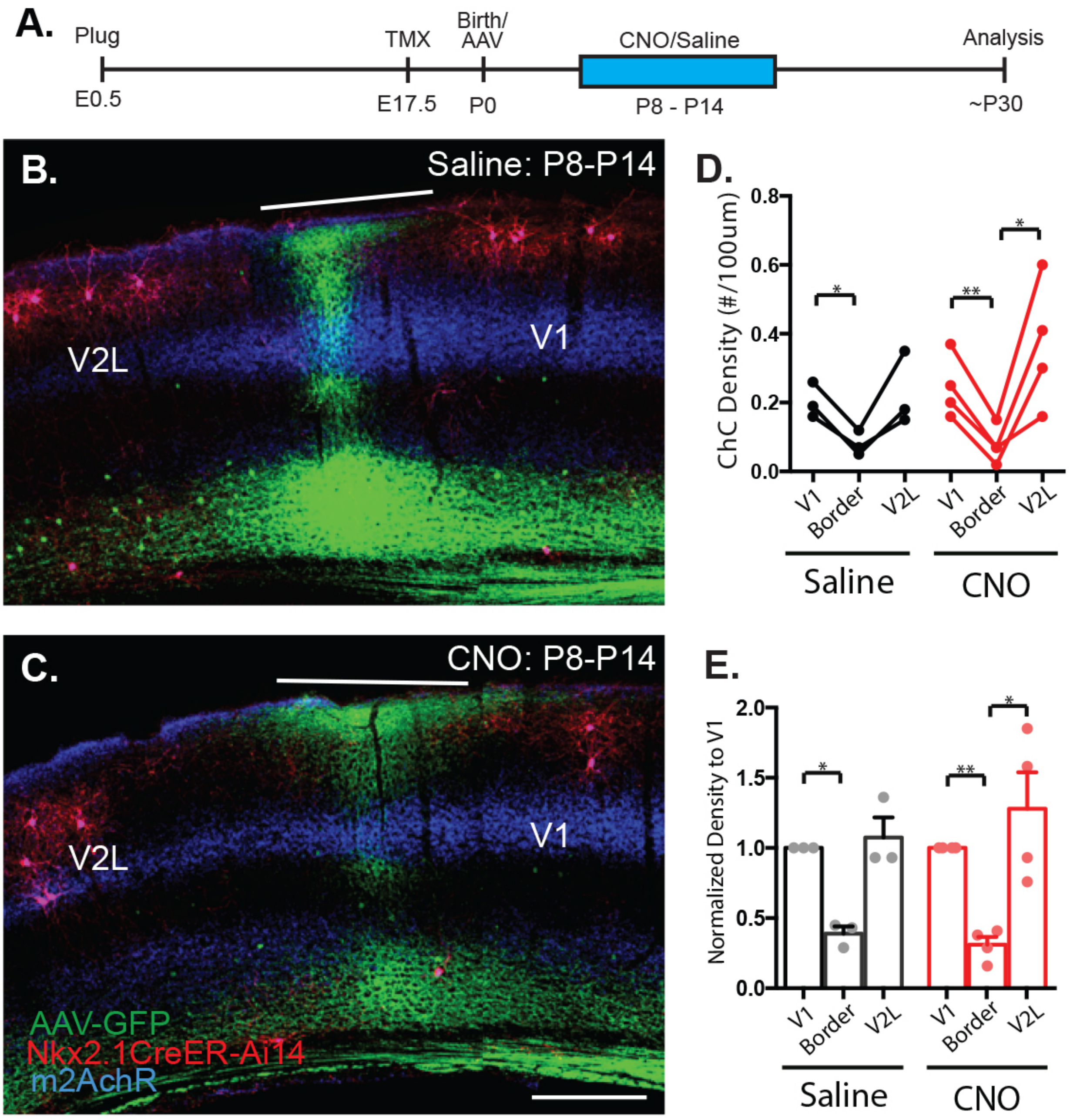


**Fig. S7. CNO does not have off-target effects on Chandelier cell elimination**

(**A**) Time course of AAV-EGFP injection at P0, tamoxifen induction at E17.5, and daily CNO or Saline i.p. injections between P8-P14. (**B-C**) Example image of ChC distribution at the border region between V1 and V2L in saline control (n = 3 animals) (**B**) and CNO-treated (n = 3 animals) (**C**) animals. (**D**) Decrease in ChC density at V1/V2L border in both saline control and CNO-treated animals. (**E**) Normalized ChC density to V1 showing CNO between P8-P14 does not have off-target effects on ChC elimination. * $p < 0.05$, ** $p < 0.01$, *** $p < 0.001$. All statistics performed with paired t-test unless stated otherwise. Scale Bar: 200um.

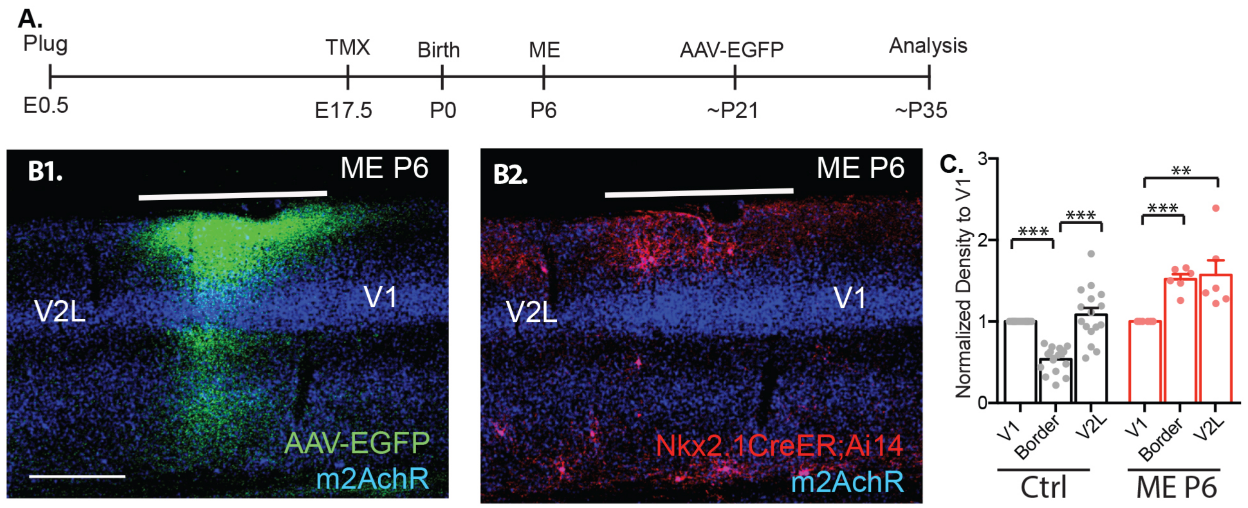


**Fig. S8. Monocular enucleation at P6 prevents Chandelier cell elimination at V1/V2L border.**

(**A**) Time course of tamoxifen induction at E17.5, monocular enucleation (ME) at P6 and AAV-EGFP at ~P21. (**B**) Example image of callosal projection (**B1**) and ChC distribution (**B2**) at the border region between V1 and V2L in ME animals (n = 6 animals). (**C**) Normalized ChC density to V1 showing ME at P6 prevents ChC elimination at the V1/V2L border. * $p < 0.05$, ** $p < 0.01$, *** $p < 0.001$. All statistics performed with paired t-test unless stated otherwise. Scale Bar: 200um.

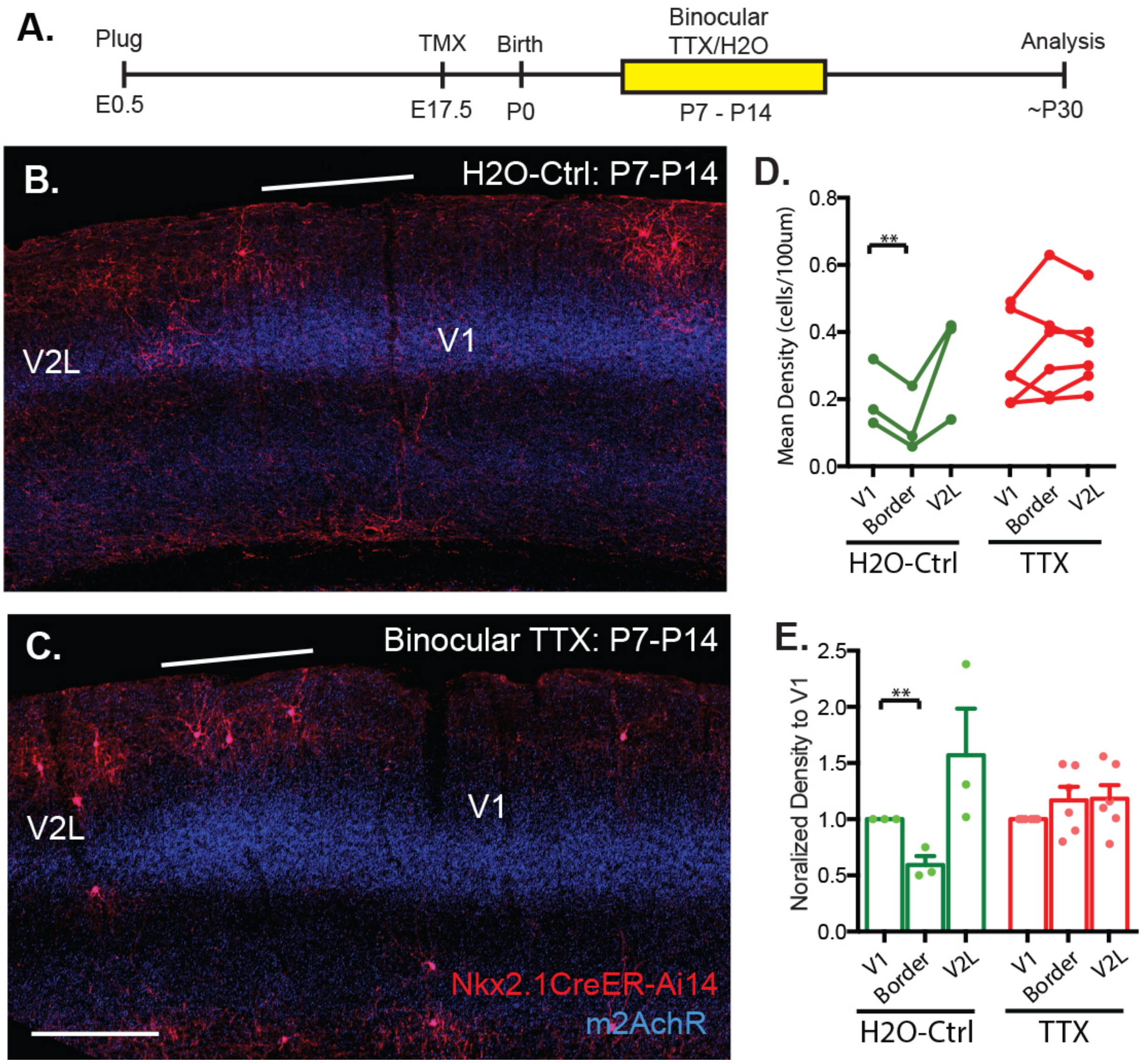


**Fig. S9. Binocular blockade of retinal activity between P7-P14 prevents Chandelier cell elimination at V1/V2L border.**

(**A**) Time course of tamoxifen induction at E17.5, and daily binocular TTX retinal injection between P7-P14. (**B-C**) Example image of ChC distribution at the border region between V1 and V2L in H2O control (B) and TTX-injected (**C**) animals. (**D**) Decrease in ChC density at V1/V2L border in saline control (n = 3 animals), but not binocular TTX-treated animals (n =

6 animals). (**E**) Normalized ChC density to V1 showing binocular TTX injection between P7-P14 prevents ChC elimination at V1/V2L border. * $p < 0.05$, ** $p < 0.01$, *** $p < 0.001$. All statistics performed with paired t-test unless stated otherwise. Scale Bar: 200um.

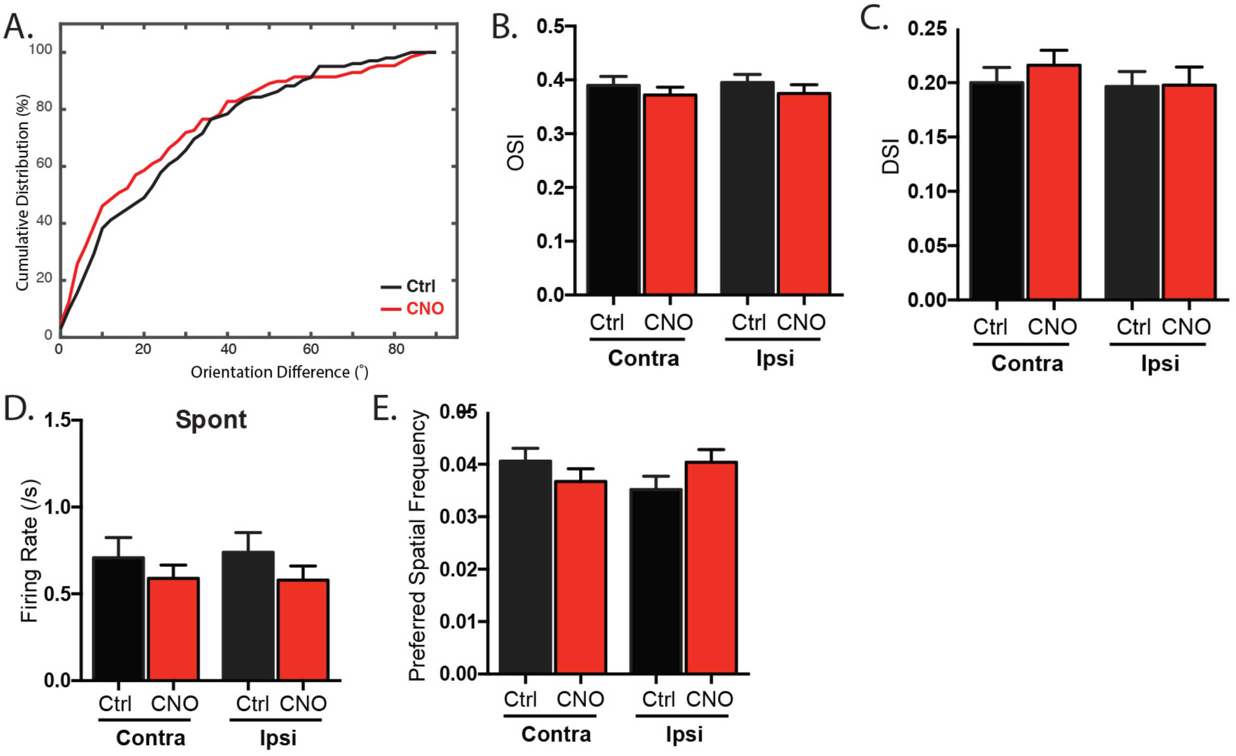


**Fig. S10. Excess Chandelier cells do not affect the development of binocular matching of orientation preference and most of monocular response properties.**

(**A**) Cumulative distribution of ΔO for control (n = 102 cells, 5 animals) and CNO treated (n = 128 cells, 7 animals) animals showing no significant difference between the two groups (p = 0.16, K-S test). (**B-D**) Mean OSI (**B**), DSI (**C**), and spontaneous firing rate (**D**) is similar in both groups for both eyes (unpaired t-test). (**E**) CNO treated and control animals have similar spatial frequency preference for both eyes (K-S test).

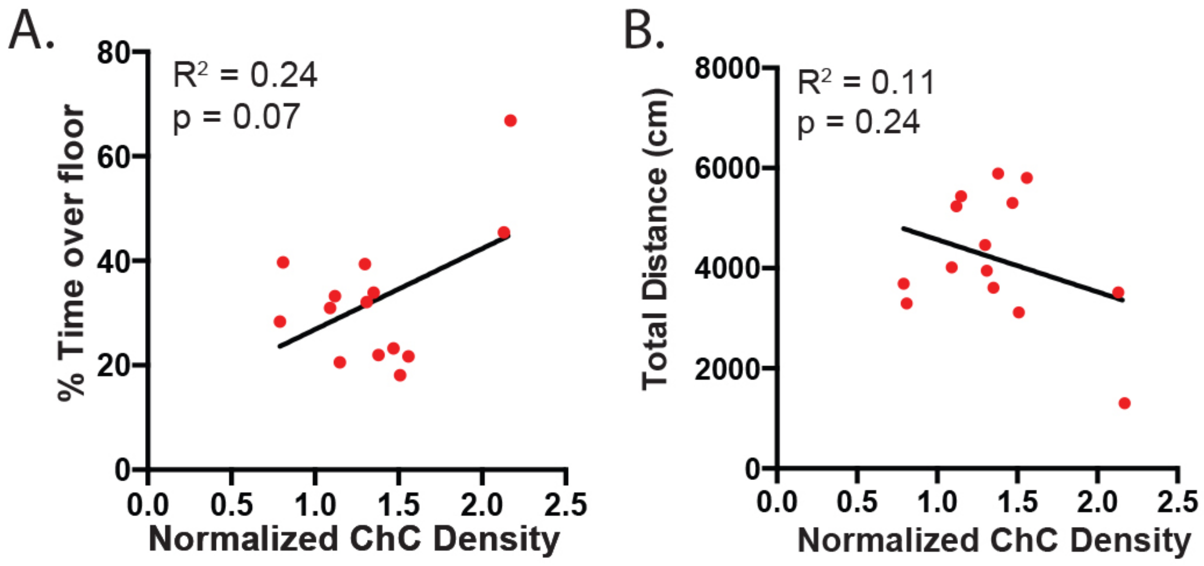


**Fig. S11. Correlation of visual cliff test parameter and Chandelier cell density at V1/V2L border.**

(**A**) Correlation of % time animal spend over the floor side and ChC density at V1/V2L border ($r$ = 0.49, p = 0.07, n = 14 animals). (**B**) Correlation of total distance the animal travelled and ChC density at V1/V2L border ($r$ = 0.33, p = 0.24, n = 14 animals).

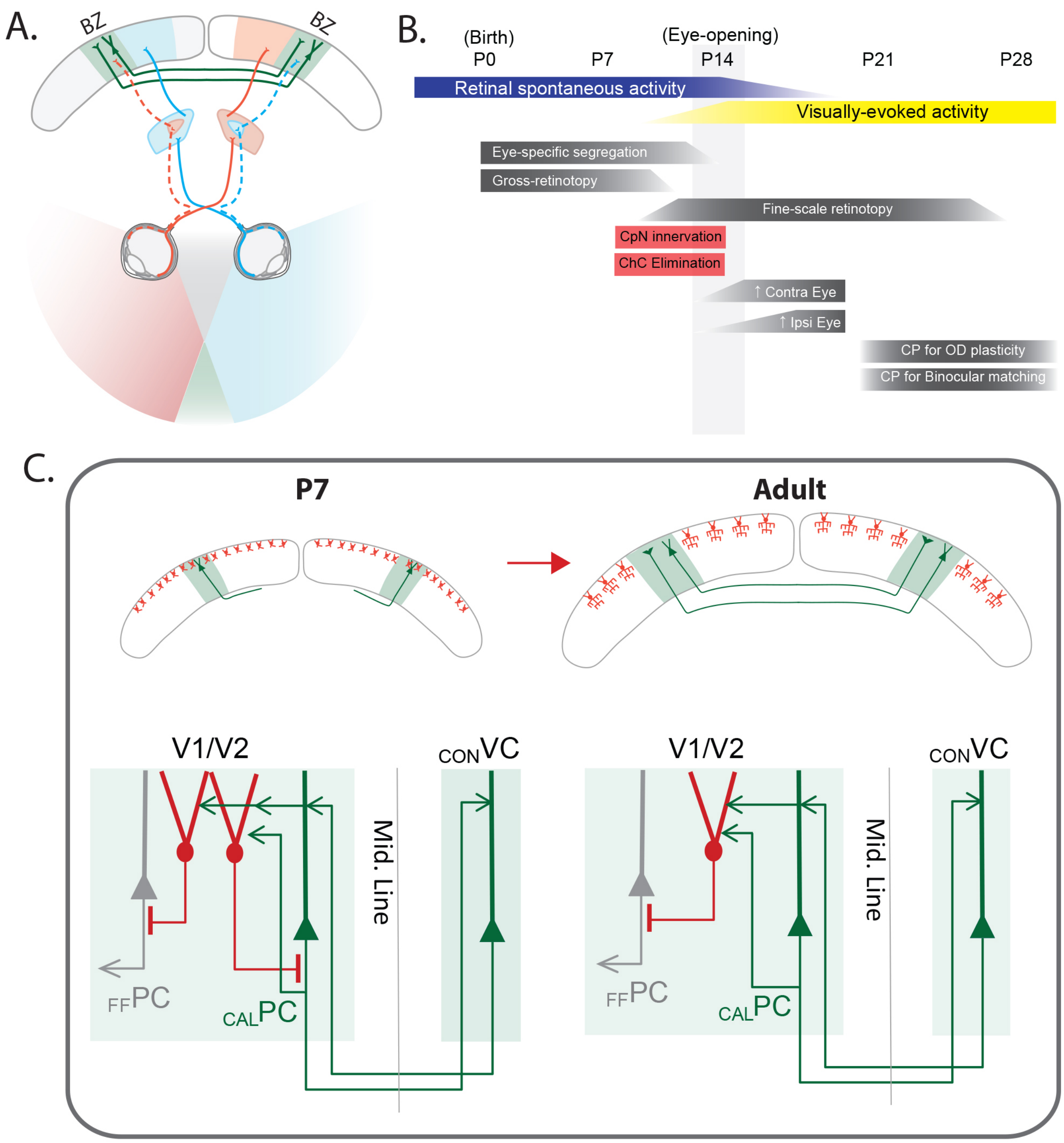


**Fig. S12. A model for Chandelier cell elimination and visual system development.**

(**A**) In rodents, information in the central visual field is first projected separately to the temporal region of the left and right retina, relayed largely in parallel through the lateral geniculate nuclei (LGN), and converge at the lateral region of the primary visual cortex (V1), defining the binocular zone (BZ, shown in green). In addition to this convergence of ipsi- and contra- lateral LGN-V1 inputs, pyramidal neurons at BZ project to the contralateral BZ, forming a transcallosal pathway that also contributes to binocular response properties. (**B**) Chandelier cell elimination takes place between P7 to P14 before eye opening. The timing coincides with axonal projections of transcallosal neurons onto the contralateral hemisphere as well as spontaneous retinal activity before eye opening (43). (**C**) At P7, abundance of ChCs at the V1/V2L border may form promiscuous connectivity between ChCs and callosal projecting neurons. As GABAergic transmission may be depolarizing at early postnatal ages, ChCs innervating callosal PyNs might promote their firing, forming a trans-callosal loop, which is further driven by coordinated bilateral retinal inputs. Such a transient over-excited network may drive the elimination of "mis-wired" ChCs through apoptosis, resulting a reduction in ChCs by P14.

**Movie S1. Example behavior of control mouse while performing visual cliff test.** The mouse is allowed to move freely around a box with a clear acrylic base. A contrast grating is placed below the box at two different heights. The animal pauses when it reaches the border between the shallow and deep halves, appears to inspect the border region but does not cross over, then retreats to the shallow end.

**Movie S2. Example behavior of CNO-treated mouse while performing visual cliff test.** Unlike the control mouse, when CNO-treated mouse approaches the border region, it crosses onto the deep side with no hesitation.
